# Structure and function of *N*-acetylglucosamine kinase illuminates the catalytic mechanism of ROK kinases

**DOI:** 10.1101/2021.09.30.462564

**Authors:** Sumita Roy, Mirella Vivoli Vega, Jessica R. Ames, Nicole Britten, Amy Kent, Kim Evans, Michail N. Isupov, Nicholas J. Harmer, on behalf of the GoVV consortium

**Affiliations:** Living Systems Institute, Stocker Road, Exeter EX4 4QD, U.K; Henry Wellcome Building for Biocatalysis, Biosciences, Stocker Road, Exeter EX4 4QD, U.K; School of Biochemistry, University of Bristol, Bristol BS8 1TD, U.K; H.H. Wills Physics Building, School of Physics, University of Bristol, Tyndall Avenue, Bristol BS8 1TL, U.K

**Keywords:** carbohydrate kinase, enzyme mechanism, magnesium, differential scanning fluorimetry, X-ray crystallography

## Abstract

*N*-acetyl-D-glucosamine (GlcNAc) is a major component of bacterial cell walls. Many organisms recycle GlcNAc from the cell wall or metabolise environmental GlcNAc. The first step in GlcNAc metabolism is phosphorylation to GlcNAc-6-phosphate. In bacteria, the ROK family kinase NagK performs this activity. Although ROK kinases have been studied extensively, no ternary complex showing the two substrates has yet been observed. Here, we solved the structure of NagK from the human pathogen *Plesiomonas shigelloides* in complex with GlcNAc and the ATP analogue AMP-PNP. Surprisingly, *Ps*NagK showed two conformational changes associated with the binding of each substrate. Consistent with this, the enzyme showed a sequential random enzyme mechanism. This indicates that the enzyme acts as a coordinated unit responding to each interaction. Molecular dynamics modelling of catalytic ion binding confirmed the location of the essential catalytic metal. Site-directed mutagenesis confirmed the catalytic base, and that the metal coordinating residue is essential. Together, this study provides the most comprehensive insight into the activity of a ROK kinase.

## Introduction

*N*-acetylglucosamine (GlcNAc) is a critical monosaccharide for both prokaryotes and eukaryotes. Eukaryotes widely employ GlcNAc in the *N*- and *O*-linked glycans that decorate protein surfaces; in the glycosaminoglycans hyaluronan, heparin sulfate and keratan sulfate that form a major part of the connective tissues (1) (2); and in chitin (3). GlcNAc is also used as a reversible modification of proteins (4) that is conserved amongst metazoans, and to decorate some growth factors (5). This modification is particularly common on nuclear proteins, and generally acts to modulate signalling (often in competition with phosphorylation) and transcription in response to stress and nutrient conditions (6–8).

GlcNAc is essential to most prokaryotes, as the cell wall is formed from a polymer of GlcNAc and *N-* acetylmuramic acid cross-linked with peptides (9). Consequently, the key enzymes required for the biosynthesis of the nucleotide linked sugar UDP-GlcNAc are essential in all bacteria. Many bacteria also require GlcNAc to form their lipopolysaccharides (with GlcNAc forming the core of lipid A) (10) and capsular polysaccharides (CPS) (11). Many oligosaccharides are initiated by the addition of GlcNAc, its epimer *N-*acetylgalactosamine (GalNAc), or 6-deoxy versions of these (*N-*acetyl-D-quinovosamine and *N-*acetyl-D-fucosamine respectively) to a lipid carrier (10, 12, 13). The wzx flippase that transfers oligosaccharides from the cytoplasmic leaflet of the inner membrane into the periplasm (14, 15) and the wzy O-antigen/CPS polymerase (16) have strong specificity for the membrane proximal sugar. Furthermore, most oligosaccharide transferases (17, 18) are exquisitely specific for the *N*-acetyl group, making the *N-*acetylated sugars intimately linked to the surface biology of bacteria.

GlcNAc is generally synthesised by cells from glucose (Figure 1) (19, 20). However, many organisms also have pathways for recycling GlcNAc. This is of particular importance for many bacteria that remodel their cell wall, and for intracellular bacteria that have a reduced availability of metabolic precursors in their environmental niches. Loss of the recycling pathway enzymes reduces the capacity of bacteria to remodel their cell walls (21–24). These pathways have been recognised in a wide range of human pathogens (e.g. *Escherichia coli* (22), *Pseudomonas aeruginosa* (25), *Enterobacteriaceae, Staphylococcus aureus* (26, 27), *Mycobacterium tuberculosis* (28)). Many bacteria utilise chitin as a nutrition resource, using chitinases to recycle it to GlcNAc (29, 30). They are likely to be of particular importance in pathogens derived from crustaceans and insects (e.g. *Serratia* (31) and *Vibrio* species (32)).

**Figure 1:**
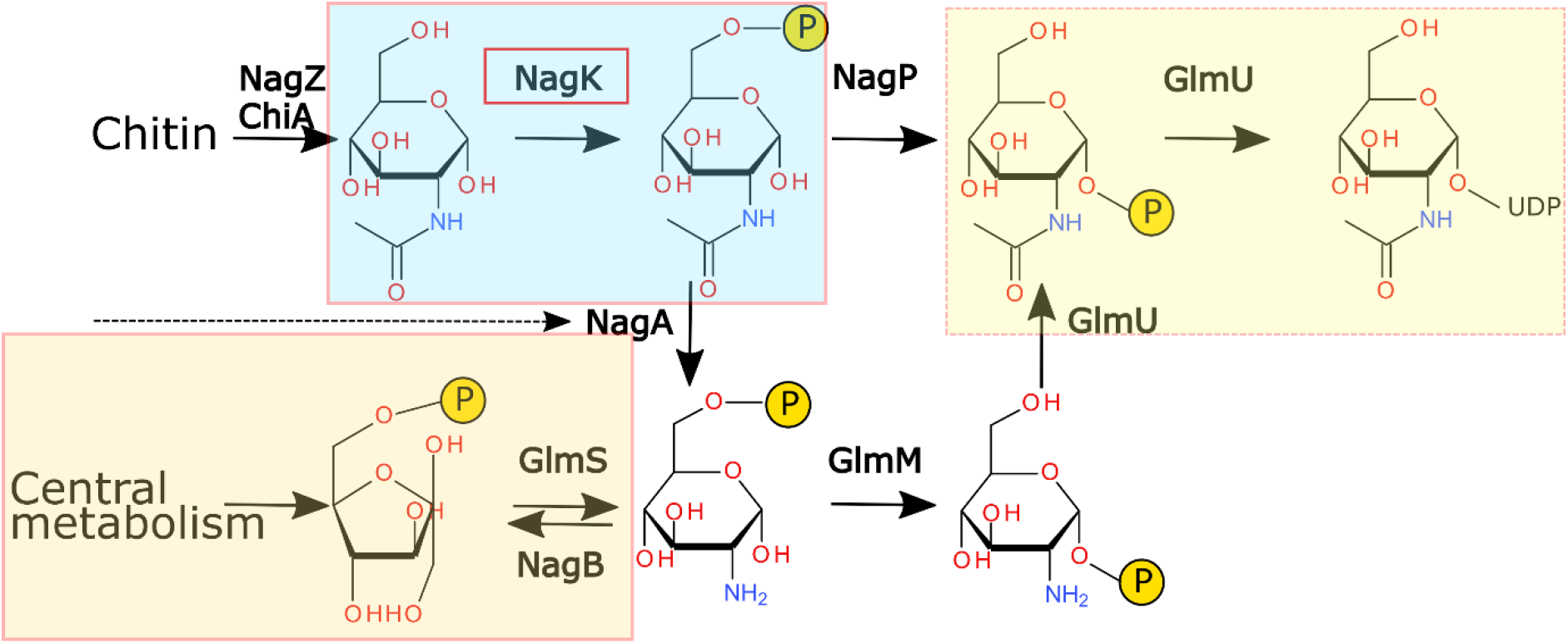
Biosynthesis and catabolism of *N*-acetylglucosamine. Cells from bacteria to humans synthesise *N-*acetylglucosamine from fructose-6-phosphate derived from the Embden–Meyerhof– Parnas pathway (orange shaded box). Some organisms utilise GlcNAc from the environment (e.g. digested chitin, bacterial cell wall components or glycosaminoglycans). GlcNAc is converted into GlcNAc-6-phosphate by NagK enzymes (sky blue shaded box). It is then either deacetylated for catabolism or recycling into cellular GlcNAc products (black dashed arrow); or isomerised to GlcNAc-1-phosphate for direct transfer to UDP (yellow shaded box).

An essential step in GlcNAc metabolism is the phosphorylation of GlcNAc to GlcNAc-6-phosphate (GlcNAc-6P). Eukaryotes isomerise this to GlcNAc-1-phosphate (33, 34) (Figure 1), as their preferred metabolic route to UDP-GlcNAc. In contrast, bacteria that recycle GlcNAc deacetylate GlcNAc-6P, linking recycled and environmental GlcNAc to their central metabolism (35). Phosphorylation of GlcNAc to GlcNAc-6P is performed by a specific kinase, *N-*acetylglucosamine kinase (NagK). Both mammalian (36) and bacterial NagK enzymes belong to the ROK kinase family of carbohydrate kinases (37). This family phosphorylates a broad range of sugars, with individual kinases showing tight specificity for their substrates (38–41). ROK kinases have a two-domain fold, with the sugar binding between the two domains, causing a structural re-arrangement that forms the active site (42, 43). Other characterised ROK kinases have shown a requirement for either manganese or magnesium for catalysis (40, 44, 45). Existing crystal structures suggest that ROK kinases use a similar mechanism to other classes of carbohydrate kinases (37) (Figure 1b). A conserved aspartic acid side chain deprotonates the 6’-hydroxyl of GlcNAc. This hydroxyl attacks the ATP γ-phosphate, passing through a pentacoordinate transition state that is stabilised by the catalytic metal. However, current structural information does not include a structure of an ATP analogue with an intact γ-phosphate. There is only one structure (from the human *N-*acetylmannosamine kinase NanK) that contains a catalytic metal: the metal binding site has not been confirmed by mutations or in bacterial enzymes (36, 46).

Here, we report the activity, structure, and mechanism of NagK from *Plesiomonas shigelloides*. Surprisingly, the enzyme displays a random sequential mechanism, with both GlcNAc and ATP able to bind to the enzyme first. PsNagK showed activity with magnesium and manganese as divalent cofactors. The structure of PsNagK in complex with GlcNAc and the ATP analogue AMP-PNP demonstrates how the enzyme catalyses phosphorylation of GlcNAc. Molecular dynamics simulations allowed us to confirm the location of the catalytic cation binding site. Comparing the ternary complex to the product complex of NagK bound to GlcNAc-6P highlights a possible catalytic mechanism. This provides, for the first time, a comprehensive kinetic and structural characterisation of a ROK kinase.

## Results

### NagK activity from divergent species

The enzymatic activity of NagK has previously been described for *E. coli* (47). We determined the activity for a wider range of enzymes, to highlight the diversity in activity from different species. We particularly focused on human pathogens with diverse NagK sequences. Recombinant NagK was readily purified for a range of human pathogens (Figures S1 and S2). The enzymes showed a range of activities (Table 1), with NagK from *Vibrio vulnificus* showing the highest activity.

**Table 1:** Activity of NagK orthologues from diverse bacteria. NagK orthologues from diverse bacteria were purified (Figure S1) and their activity determined. Apparent *K*_*M*_ values were determined in the presence of at least 10X *K*_*M*_ of the partner substrate. The sequences were aligned using MUSCLE and percentage identities calculated in Geneious v. 8.

| Species | $k_{cat}$ ($s^{-1}$) | $K_{M\ app\ GlcNAc}$ ($\mu M$) | $K_{M\ app\ ATP}$ ($\mu M$) | Identity to *E. coli* NagK (%) |
| --- | --- | --- | --- | --- |
| *Vibrio vulnificus* | 95 ± 8 | 209 ± 56 | 41 ± 9 | 54.5 |
| *Photobacterium damsela* | 230 ± 10 | 60 ± 10 | 250 ± 10 | 53.8 |
| *Pseudoalteromonas* sp. P1-8 | 59 ± 7 | 1100 ± 200 | 3000 ± 700 | 40 |
| *Plesiomonas shigelloides* | 202 ± 3 | 98 ± 6 | 290 ± 20 | 59.7 |

### NagK uses a sequential mechanism

We selected NagK from *P. shigelloides* for a more detailed study of the NagK mechanism. The enzyme kinetics showed a sequential mechanism rather than a ping pong mechanism (Figure 2A-C; *p*=0.0045). The products GlcNAc-6P and ADP showed weak inhibition, with Morrison *K*_*i*_ values one to two orders of magnitude higher than the cognate substrate *K*_*M*_ (Figure S3). This prevented determination of whether an ordered or random sequential mechanism is used as alternative interpretations would be within error. We therefore examined whether the binding of either substrate affects binding of the other. Past studies of enzyme mechanisms have investigated substrate binding using methods such as differential scanning fluorimetry (48) or fluorescence anisotropy (49). We used differential scanning fluorimetry to determine the dissociation constants of GlcNAc and the non-hydrolysable ATP analogue AMP-PNP. Using the isothermal DSF approach (50), we determined that the *K*_*D*_ for GlcNAc in the absence and presence of AMP-PNP were 230 ± 20 μM and 270 ± 20 μM respectively (Figure 2D). We chose a temperature of 68 °C to measure at as this gave the optimal signal to determine *K*_*D*_. The *K*_*D*_ for AMP-PNP in the absence and presence of GlcNAc were 2.2 ± 0.6 mM and 3.2 ± 0.5 mM respectively (Figure 2E). Surprisingly, there is no significant increase in the affinity for either substrate in the presence of the other. This suggests that NagK uses a random sequential mechanism.

**Figure 2:**
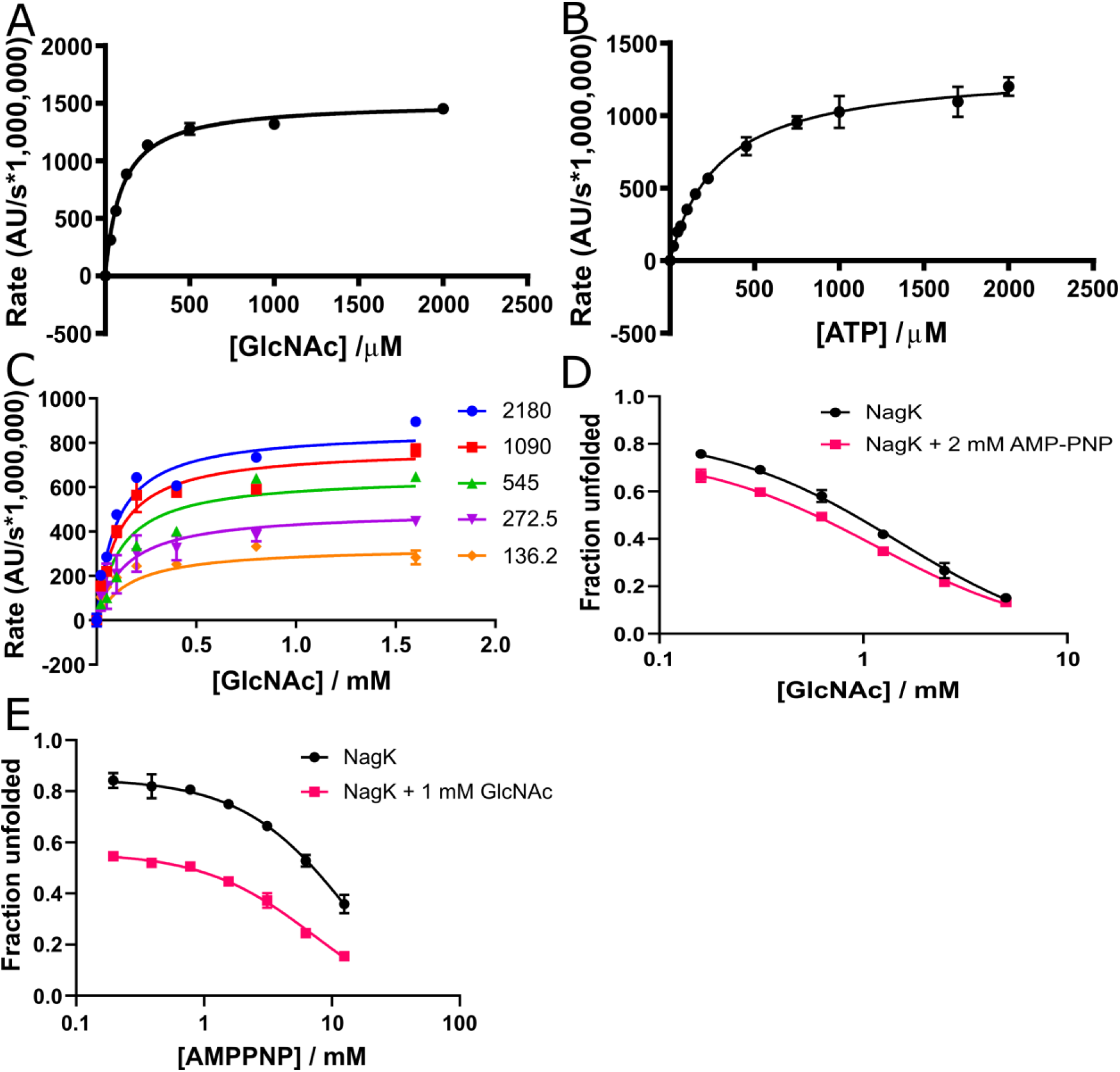
NagK uses a sequential mechanism of binding. NagK from *Plesiomonas shigelloides* was selected for a more detailed study of function as it has intermediate activity. PsNagK activity fitted to the Michaelis-Menten equation for both GlcNAc (**A**) and ATP (**B**). (**C**) Testing of both substrates together showed a strong preference to the equation for sequential binding rather than a ping-pong mechanism (Akaike’s information criteria difference = 10.79; *p* = 0.0045). Neither GlcNAc-6-P nor ADP showed product inhibition at readily testable concentrations (Fig S3), preventing determination of whether the binding is ordered or random. (**D, E**) Differential scanning fluorimetry of NagK in the presence of GlcNAc and the ATP analogue AMP-PNP. The apparent *K*_*D*_ value of NagK for GlcNAc shows no significant difference in the absence (230 ± 20 μM) or presence (270 ± 20 μM) of 2 mM AMP-PNP. The apparent *K*_*D*_ value of NagK for AMP-PNP increases slightly from 2.2 ± 0.6 mM to 3.2 ± 0.5 mM in the presence of 1 mM GlcNAc.

### NagK prefers magnesium as the catalytic metal

Most carbohydrate kinases require a metal cofactor. The ROK kinases particularly have previously shown a strong requirement for metals. Consistent with this, *Ps*NagK showed no activity in the absence of divalent cations (Table 2). The enzyme showed a preference for manganese (*K*_1/2_ = 0.07 ± 0.01 mM) over magnesium (*K*_1/2_ = 0.32 ± 0.05 mM) at low concentrations, but at higher concentrations manganese was inhibitory (*K*_i_ = 11 ± 2 mM; maximum rate 55 ± 3 s^−1^ at 0.87 mM; Figure 3A). Magnesium shows no inhibition and a higher maximum rate (102 ± 3 s^−1^) and would be strongly preferred at physiological concentrations (Figure 3B). No activity was observed with calcium.

**Table 2:** Effect of divalent cations on enzyme activity. *P. shigelloides* NagK was tested at *K*_*M*_ for both substrates. Data for magnesium were fitted to the Michaelis-Menten equation. Data for manganese were fitted to the substrate inhibition equation. Data were fitted in GraphPad v. 8.0.

| Metal | Maximum rate ($s^{-1}$) | $K_{1/2}$ (mM) | $K_i$ (mM) |
| --- | --- | --- | --- |
| Mg<sup>2+</sup> | 105 ± 3 | 0.31 ± 05 | N/A |
| Mn<sup>2+</sup> | 65 ± 3 | 0.07 ± 01 | 11.3 ± 2.4 |
| Ca<sup>2+</sup> | 0 | N/A | N/A |

**Figure 3:**
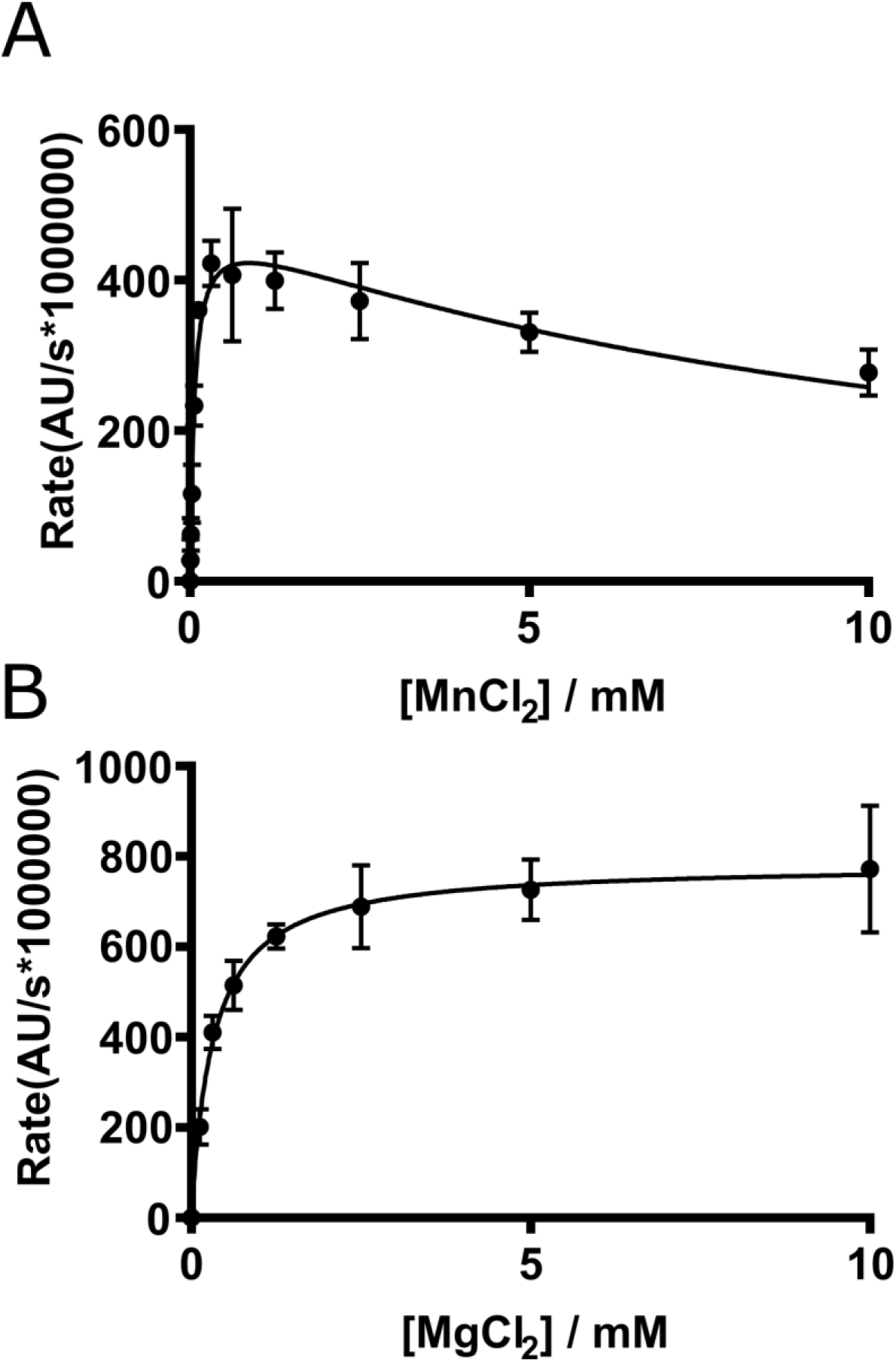
Magnesium is the preferred metal cofactor of NagK. The activity of *P. shigelloides* NagK was tested in the presence of magnesium, manganese, or calcium. No activity was observed without divalent cations or with calcium. Manganese (**A**) shows a strong activity at low concentrations (maximum rate 55 ± 3 s^−1^ at 0.87 mM), but higher concentrations inhibit activity (*K*_*I*_ = 11 ± 2 mM). Magnesium (**B**) shows a maximum rate of 102 ± 3 s^−1^, with *K*_*1/2*_ = 0.32 ± 0.05 mM, and shows no evidence of inhibition.

### The NagK active site is formed by enzyme closure around the GlcNAc and ATP substrates

Although a structure of *V. vulnificus* NagK has been solved (51), there is no structure of a ligand bound NagK. We therefore determined the structure of *P. shigelloides* NagK, as this crystallised readily with and without its substrates (Table S1). As expected, *Ps*NagK forms a two-domain fold with a large domain (including the structural zinc characteristic of ROK kinases (37)) and a small domain (Figure 4A). The enzyme closes around the GlcNAc substrate, with the small domain rotating by 23° (moving up to 15 Å) relative to the large domain (Figure 4B). The GlcNAc is bound specifically by the side chains of residues S78, N104, D105, E154, H157, and D187 (Figure 4C).

**Figure 4:**
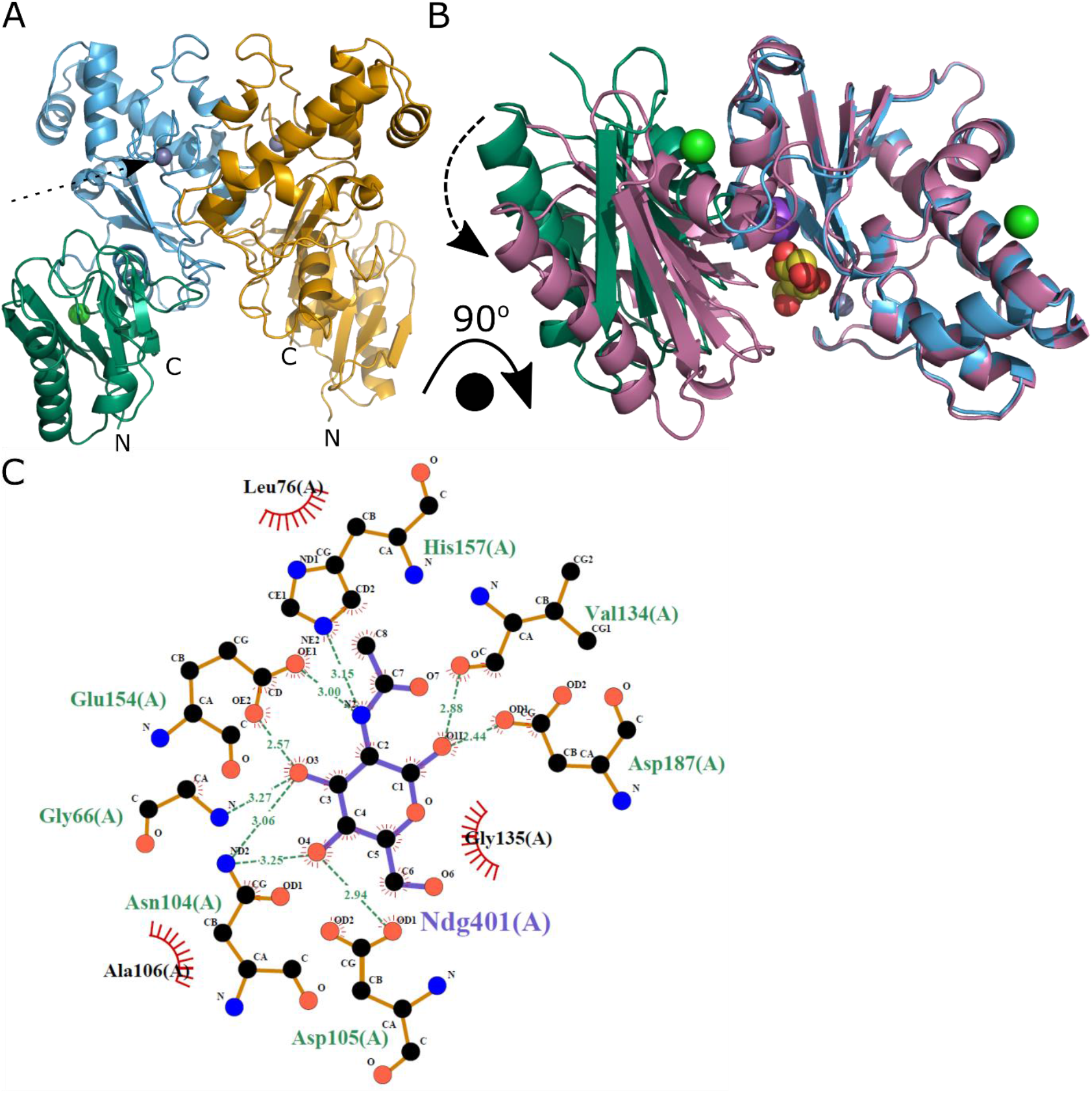
NagK closes around the *N*-acetylglucosamine substrate. **A:** Overall structure of the NagK dimer. NagK has two domains: an N-terminal small domain (green) that includes the C-terminal helix, and a large domain (sky blue). NagK forms a dimer (second molecule in yellow), with the interface between two large domains. A conserved structural zinc ion (grey sphere) is seen in the large domain (dashed arrow). The N and C-termini are indicated. **B:** Upon binding of the ligand *N-*acetylglucosamine (spheres, carbon atoms in yellow), the small domain rotates approximately 15° relative to the large domain to close around the sugar. Unbound structure coloured as in **A**; bound structure shown in magenta. Structures were superimposed over the large domain. **C:** GlcNAc is held in place by a network of amino acids from both small (residues 1–104, 291–303) and large (residues 105-290) domains. The interactions give the enzyme good specificity for GlcNAc over possible competing substrates. Structure images shown as cartoon with ligand atoms shown as spheres. Atom colours where not indicated: nitrogen, blue; oxygen, yellow; chloride, green, potassium, purple. Panels A and B generated using PyMOL v. 2.4.1 (27); panel C generated using LigPlot+ v2.2 (81, 82).

We then soaked the ATP analogue AMP-PNP into the *Ps*NagK structure. A structure with both GlcNAc and AMP-PNP shows the location of the γ-phosphate in a position poised for catalysis (Figure 5A): the best previous ROK kinase ligand structures showed density only to the β-phosphate (42, 46). The small domain rotates a further 16° to engage the ATP (Figure S4). ATP is held in place by the side chains of residues T10, D105, T132 and E196, with the phosphates being coordinated by the main chain of G9, T10, and G255 (Figure 5B). Most of these side chains are well conserved amongst NagKs, consistent with a role in substrate binding (Figure S5). We were unable to obtain a structure that contained the catalytic cation. However, our ternary complex with GlcNAc and AMP-PNP is structurally very similar to the previously solved NanK structure that included a catalytic magnesium ((46); Figure S6). The cation binding site is adjacent to a water molecule in our structure coordinated by D6, the main chain carbonyl of I7, and the γ-phosphate (Figure 5C). To test the hypothesis that this is the metal binding site, we performed molecular dynamics simulations of the active site with divalent cations added in this location, and AMP-PNP replaced by ATP. Molecular dynamics of the solved structure over 5 ns showed no significant changes in the structure, aside from a minor re-arrangement of the ATP phosphates (Figure S7A). When magnesium, manganese or calcium was added to the protein structure, the cation and ATP phosphates re-arrange to form a binding site for the divalent cation. Counterintuitively, in the cases of magnesium and manganese, the re-arrangement brings the cation close to the side chain of D105 and the GlcNAc O6 as well as the D6 side chain, I7 main chain carbonyl and the γ-phosphate (Figure S7B, D). These cations show pentahedral coordination as one face is partially blocked by the side chain of I127. In contrast, the calcium ion forms a classical octahedral coordination with the side chains of D105 and D6 (both oxygens), I7 main chain, and two oxygens from the ATP γ-phosphate. In this case GlcNAc O6 is excluded from the coordination. This may reduce the acidity of the GlcNAc O6, consistent with calcium not supporting catalysis. The rapid, reproducible re-arrangement of the active site under molecule dynamics strongly supports the hypothesis that this is the cation binding site. The cation is then positioned to stabilise the pentacoordinate transition state. However, it is likely that a further re-arrangement of the enzyme active site is necessary for catalysis, as the ATP γ-phosphate remains too far away from GlcNAc to support a reaction.

**Figure 5:**
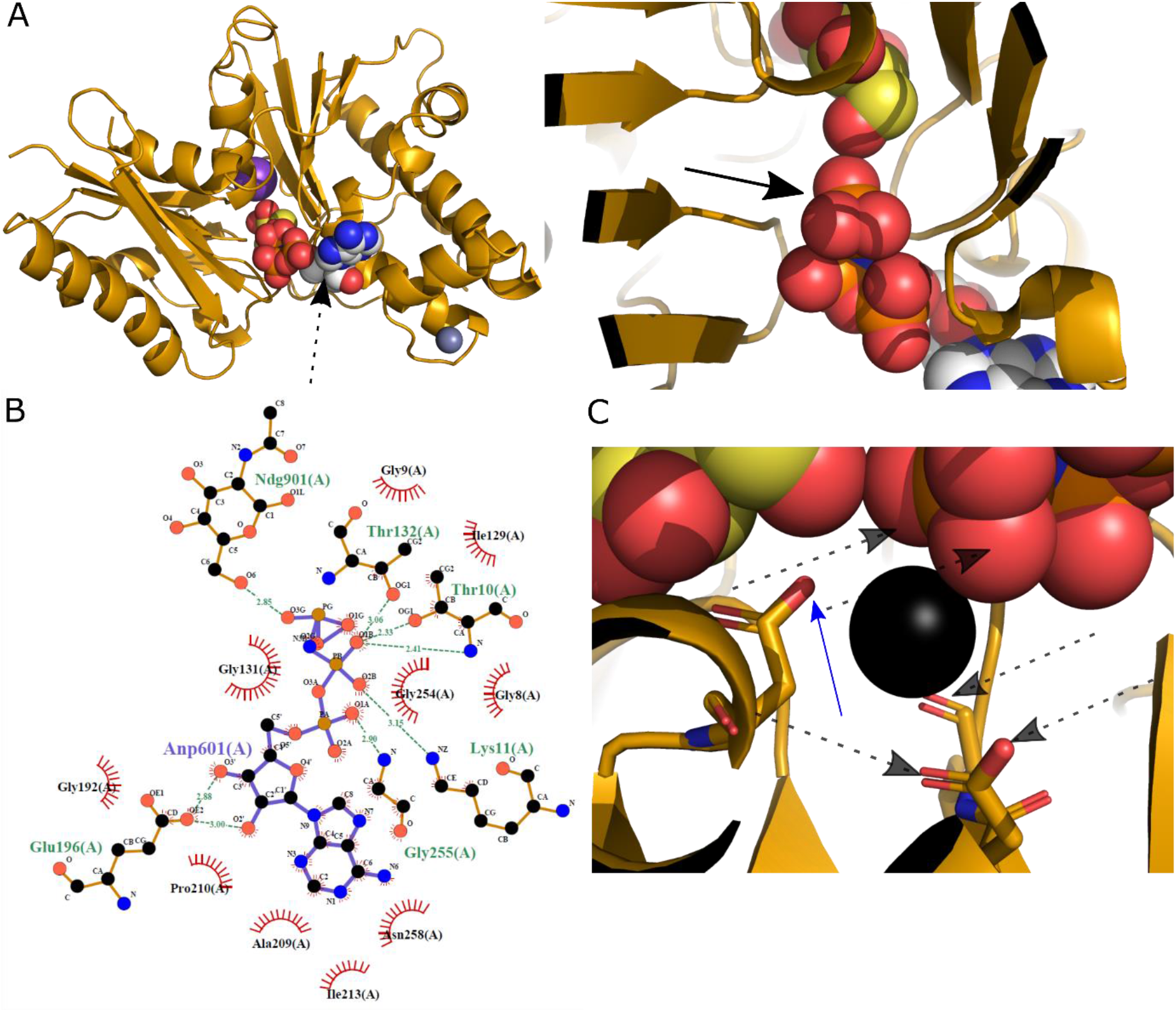
The NagK ternary complex with GlcNAc and ATP highlights the likely catalytic mechanism. **A:** Structure of the NagK-GlcNAc-AMP-PNP ternary complex. Left: overview of the structure in the same conformation as figure 4B. The AMP-PNP (black dashed arrow) is shown as spheres with carbon atoms coloured white. Right: close-up view of the interaction between the two ligands. The terminal phosphate is indicated with the black arrow. **B:** AMP-PNP is held in place by hydrophobic contacts to the adenine ring, hydrogen bonds from the ribose ring to E196, and interactions of the phosphate groups with T10, K11 and the protein main chain. **C:** The ternary complex creates a metal binding site that is occupied by water in the structure. Likely water molecule shown as a black sphere. Likely metal coordinating atoms (acid oxygens of E6, main chain carbonyl of I7, γ-phosphate atoms) are indicated with grey dashed arrows; the side chain of D105 (blue arrow) may act as a sixth ligand. Atom colours where not indicated: nitrogen, blue; oxygen, yellow; chloride, green, potassium, purple; zinc, grey; phosphorus, orange. Panels A and C generated using PyMOL v. 2.4.1 (27); panel B generated using LigPlot+ v2.2 (81, 82).

### Confirmation of proposed ligand interacting residues by site-directed mutagenesis

Site-directed mutagenesis of proposed ligand binding and catalytic residues support the role of these amino acids in *Ps*NagK activity. Mutation to either D105N (catalytic base) or D6N/A (metal coordinating negatively charged group) results in a loss of activity below the limit of detection (at least 1000-fold; Table 3). Mutation of the phosphate coordinating T10V and T132V results in a loss of activity, without substantially affecting the *K*_*M*_ for either substrate. Mutation of the main ATP binding side chain D187N results in an increase in *K*_*M*_, but little impact on rate. Mutation of some side chains that coordinate GlcNAc (N104D, E154Q or double mutant) results in substantial increases in *K*_*M*_ for both substrates, and clearly reduced rates. Mutation of other conserved GlcNAc binding residues S78A and E196Q resulted in clear increases in rate without affecting *K*_*M*_. These two residues are not well conserved (Figure S5), and the residues mutated to are found in other orthologues. We did not mutate H157 as this residue also coordinates to the structural zinc atom, and mutation would likely significantly affect the protein structure.

**Table 3:**
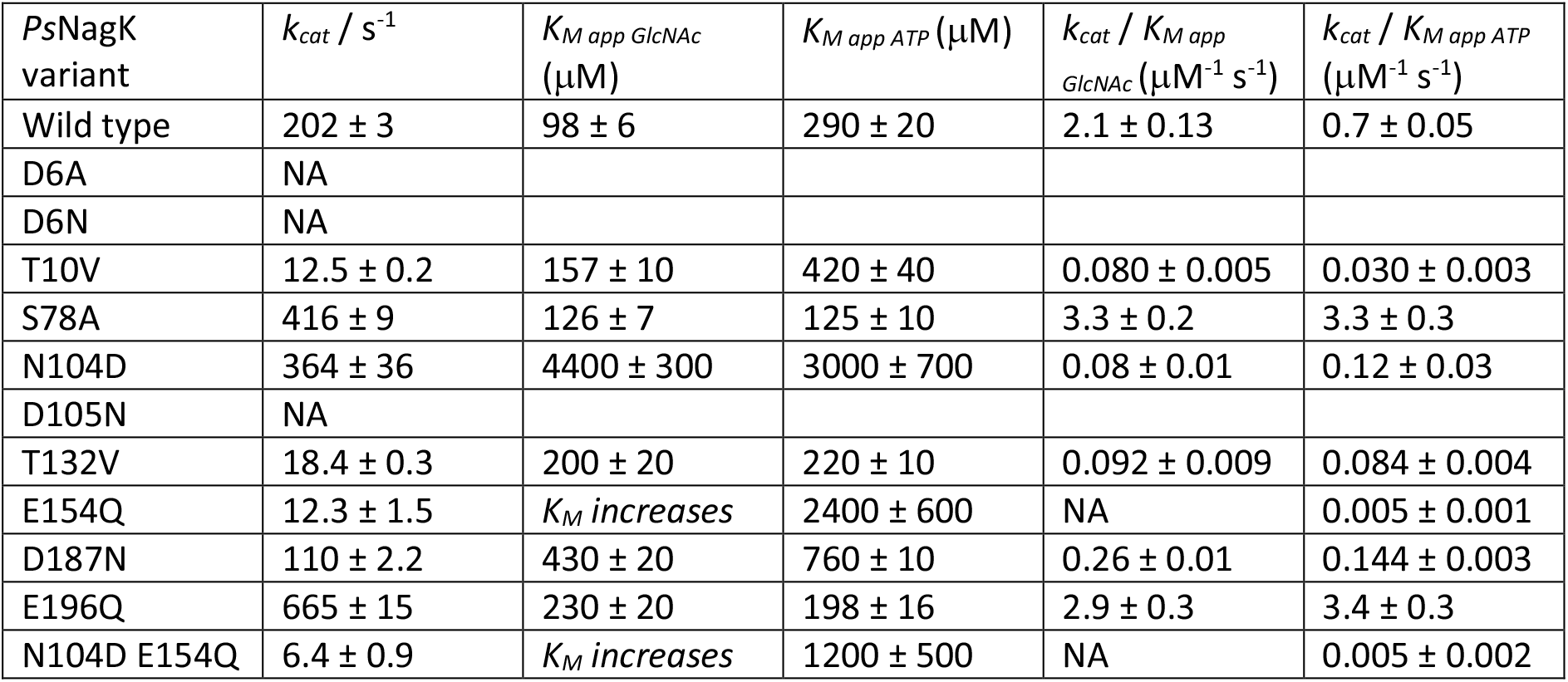
Effect of mutants of ATP and GlcNAc binding residues. Site-directed mutants were prepared for *P. shigelloides* NagK at key side chains that coordinate ATP, GlcNAc or magnesium. *k*_*cat*_ and *K*_*M app*_ for both substrates were determined as for the wild-type enzyme. D6A, D6N and D105N mutants caused a loss of activity below the limit of detection of the assay (*k*_*cat*_ < 0.01 s^−1^). The E154Q and N104D/E154Q mutants caused the apparent *K*_*M*_ for GlcNAc to increase to above 50 mM (i.e., the plot of rate against substrate concentration was a straight line).

## Discussion

GlcNAc recycling from the cell wall is important for the biology of many human pathogens. These include some of the ESKAPE pathogens (52) of greatest concern for antimicrobial resistance (22, 25–27). To efficiently recycle cell wall GlcNAc, bacteria phosphorylate and then de-acetylate GlcNAc to form glucosamine-6-phosphate (35), an intermediate in the essential UDP-GlcNAc biosynthesis pathway (Figure 1; (19, 20)). Here, we have thoroughly characterised the first enzyme that performs the first of these steps, NagK. This enzyme belongs to the ROK kinase family of carbohydrate kinases (37). Key questions arising from previous studies of ROK kinases were the order of binding of substrates; confirming the location of the catalytic metal ion; and the location of the γ-phosphate.

In common with previous ROK kinases, we determined that NagK has an absolute requirement for divalent cations (40, 45, 53). Both magnesium and manganese, but not calcium, support NagK function. Physiologically, magnesium would likely be preferred as bacterial intracellular magnesium concentrations (~2 mM) exceed *K*_1/2_ (0.3 mM), whilst manganese concentrations (5-15 μM) are below *K*_1/2_ (70 μM) (54, 55). Comparison of the crystal structure of NagK bound to GlcNAc and AMP-PNP to the human NanK structure (46) suggested that the metal ion should bind into a pocket adjacent to the γ-phosphate. This pocket would be coordinated by two oxygens from the γ-phosphate, the main chain carbonyl of I7, and the side chain of D6. An alignment of ROK kinases shows that D6 is strongly conserved as an acidic residue (Figure S7). This has previously been proposed (albeit with limited evidence) as a metal ion binding residue (36). To support this proposal, we added a magnesium ion to this site in our structure and performed a molecular dynamics simulation. The maintenance of the ion in this location is strongly supported in the simulation, with both magnesium and manganese predicted to coordinate to both substrates. Furthermore, mutation of D6 to either asparagine or alanine completely abolishes the activity of the enzyme. Given that D6 is not close to either substrate in the crystal structure, this very strong phenotype strongly supports a role in binding to the catalytic metal ion. These observations strongly support this pocket as the metal binding site for a wide range of ROK kinases.

The effect of mutations in GlcNAc binding residues is in accordance with previous studies. A detailed phylogenetic study proposed that the 3’-OH is coordinated by asparagine (N104) and glutamic acid (E154) (39). Mutations in either of these residues significantly reduced the activity of NagK. In contrast, two side chains that contact GlcNAc in the crystal structures (S87 and E196) are not evolutionarily conserved (Figure S5). Mutation of these side chains increases the catalytic efficiency of NagK *in vitro*.

Our structures provide for the first time a complex of a ROK kinase poised for activity. The structure shows the ATP γ-phosphate positioned above the 6’ OH group of GlcNAc. The catalytic base, D105, is in position to de-protonate the 6’ O and turn this into a strong nucleophile. The location of the phosphate group allows coordination of two oxygens with the catalytic metal ion. Other carbohydrate kinases generally follow a mechanism of a nucleophilic substitution with a pentahedral intermediate stabilised by a metal ion (37, 56–58). Based on ours and others’ structures, it seems highly likely that ROK kinases follow a similar mechanism.

In conclusion, our study provides for the first time a detailed explanation for the catalytic power of ROK kinases. Our data show how this family of enzymes support the pentahedral intermediate required for phosphate transfer from ATP to GlcNAc. We demonstrate that a metal ion is required for NagK enzymes, and that the conserved ROK kinase metal coordinating acid is essential for enzyme activity. Our data confirm the critical side chains that support NagK substrate selectivity for GlcNAc. The availability of a detailed structure of the catalytic state of ROK kinases will enable the engineering of these enzymes to phosphorylate alternative substrates to support synthetic biology. This enzyme would also be an attractive target for the development of small molecule inhibitors to target bacteria that rely on cell wall remodelling as part of their pathogenic processes.

## Methods

### Construction of expression vectors

a codon optimised *nagK* gene from *Plesiomonas shigelloides* was cloned into the pOPINS3C expression plasmid (N-terminal polyhistidine and SUMO tags; Addgene #41115 (59)). The gene sequence was obtained as a gBlock (IDT). DNA was amplified by PCR using the following primers: forward: 5’-cagcggtctggaagttctgtttcagggtacc-3’; reverse: 5’-aagctttctagaccagtttgtgattaacctc - 3’. Each PCR reaction contained 1ng/µL of gBlock DNA, 0.25 µM each primer, 2.5 mM dNTPs, 1X Phusion buffer and 2 U Phusion Polymerase (NEB). The PCR protocol used was an initial denaturation of 30 s at 98 °C, followed by 35 cycles of 10 s at 98 °C, 10 s at 55 °C and 1 min at 72 °C, with a final elongation step of 5 min at 72 °C. The PCR fragment and the plasmid (pOPINS3C) were assembled using the NEBuilder HiFi DNA Assembly kit (NEB), following the manufacturer’s recommendations. The assembled product was transformed into 5-alpha competent cells (NEB) and the insert sequence was confirmed by Sanger sequencing (Source Bioscience). Codon optimised *nagK* genes from *P. shigelloides, Vibrio vulnificus*, *Yersinia pseudotuberculosis, Pseudoalteromonas sp. P1-8* and *Photobacterium damselae* were cloned into pNIC28-Bsa4 (N-terminal polyhistidine tag; Addgene #26103 (60)), pOPINS3C, and pGAT2 (N-terminal polyhistidine and GST tags; Addgene #112588; last three genes (61)) by Twist Bioscience. Genbank files for all plasmids are available in Supplementary Data. Plasmid were transformed into the expression strain *E. coli* BL21 (DE3) (Novagen) using ampicillin (100 μg/mL; pGAT2 and pOPINS3C clones) or kanamycin (50 μg/mL; pNIC28 clones) for selection.

### Expression and purification of NagK

NagK was expressed in 1 litre of high salt LB broth supplemented with 100 μg/mL ampicillin or 50 μg/mL kanamycin as appropriate. Each flask was inoculated with 10 mL of an overnight culture and grown at 37 °C with shaking at 200 rpm until OD_600_ reached 0.6. NagK expression was induced with 200 μM isopropyl thio-β-D-galactoside (IPTG), and cultures were grown at 20°C for 18h. Cells were harvested by centrifugation at 4500x*g* for 30 min at 4 °C. The pellet was resuspended in binding buffer (20 mM Tris-HCl, 500 mM NaCl, 10 mM imidazole, pH 8.0) and lysed by sonication (SONIC Vibra cell™ VCX130). The lysed sample was clarified by centrifugation (24 000x*g* for 30 min at 4 °C). The soluble fraction was purified using an ÄKTAxpress chromatography system (GE Healthcare). The sample was purified firstly using a 1 mL HisTrap crude column (GE Scientific). After loading sample, the column was washed with binding buffer, and the protein eluted into binding buffer with imidazole at 250 mM. The product was purified over a Superdex 200 16/60 size-exclusion column (GE Healthcare) and eluted isocratically into 10 mM HEPES, 500 mM NaCl, pH 7.5. The eluted protein was concentrated using a Vivaspin centrifugal concentrator (Generon) to 1 mg/ml and stored at −20°C with 20% (v/v) glycerol for enzymatic assays; or concentrated to 11.5 mg/ml and stored at −80°C in small aliquots without any glycerol for crystallization.

### Kinetic analysis

NagK activity was assayed using the previously described coupling reaction with pyruvate kinase (PK) and lactate dehydrogenase (LD; (62)). For *P. shigelloides*, the His-tagged protein was used. Reactions contained 90-6000 ng/mL NagK, 40 mM HEPES, pH 7.5, 100 mM KCl, 8 mM MgCl_2_, 5 mM DTT, 100 μg/ml BSA, 200 μM NADH, 500 μM phosphoenolpyruvate, 2 U/mL PK-LD (Merck #P0294), 2 mM GlcNAc and 1 mM ATP. Reactions were performed in 96 well flat-bottomed plates (Greiner #655001) in a total reaction volume of 200 μL. Reactions were monitored by measurement of the absorbance at 340 nm over 40 min in an Infinite M200PRO plate reader (Tecan) with incubation at 37 °C. Three experimental replicates were performed for all reactions.

Kinetic parameters (*K*_M_ and *k*_cat_) for ATP and GlcNAc were determined by varying either ATP or GlcNAc concentrations between 2-0.02 mM and 2-0.03 mM respectively, keeping all other parameters constant. The data were fitted to the Michaelis-Menten equation in Prism 7.05 (GraphPad). To determine the substrate mechanism, the initial reaction rates were measured with a two-fold dilution of GlcNAc from 2 mM in eight steps, and with a two-fold dilution of ATP from 2180 μM in five steps. Two experimental replicates were taken for each data point. Data were fitted to the sequential bi-bi and ping-pong equations in Prism 9.01 (GraphPad) (62–64). To determine the effect of divalent cations, initial reaction rates were determined with the MgCl_2_ in the mixture above substituted with of 10 mM MgCl_2_, MnCl_2_, CaCl_2_, CuCl_2_, or CoCl_2_, and normalized to the rate with MgCl_2_. *K*_M_ and *V*_max_ were determined for MgCl_2_ or MnCl_2_ by varying the concentration between 0-10 mM, with GlcNAc and ATP at the determined *K*_M_. Three experimental replicates were performed for all reactions.

### Differential scanning fluorimetry

The dissociation constants (*K*_*D*_) for NagK with its substrates was determined using differential scanning fluorimetry (50). Each sample contained 0.1 mg/mL NagK, 8X SYPRO Orange dye (Fisher Scientific #10338542), 10 mM Hepes pH 7.5, 100 mM KCl and varying concentrations of either GlcNAc, AMP-PNP or the combination of these in a total volume of 10 μL. Data were collected on a Rotorgene Q (Qiagen) using the ROX channel to collect data. The melt curves showed a monotonic melt. Raw data were converted to a percentage unfolded using the fluorescence readings at the start and end of the melt to define 0 and 100% unfolded. 68 °C was selected as the temperature giving an optimal range of unfolding percentages. Data were fitted to equation **1** using Graphpad v. 9.0.1.

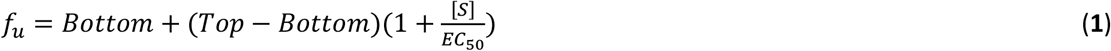

Where *f*_*u*_ is the fraction unfolded, Top and Bottom are the maximum and minimum unfolded fractions, [S] is the varied substrate concentration, and EC_50_ is the substrate concentration that reduces the unfolded fraction by half. Equations fixing Bottom as zero and including a Hill slope were rejected as inferior to **1** for these data based on Akaike’s information criterion.

The fitted EC_50_ values were converted to *K*_*D*_ using equation **2** (50).

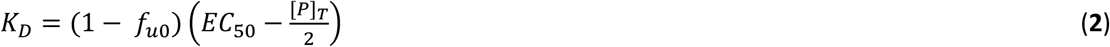

Where *f*_*u0*_ is the fraction unfolded at zero substrate concentration, and [P]_T_ is the total protein concentration.

### Crystallization

For crystallization, the His-SUMO tagged *P. shigelloides* NagK was used. Crystals were grown using the microbatch method using an Oryx8 crystallization robot (Douglas Instruments). Initial crystals grew in well E6 of the Morpheus I screen (Molecular Dimensions), mixed 1:1 with 5 mg/ml NagK. Seed stocks were prepared from these crystals in 0.1 M MOPS pH 7.5, 30% (v/v) ethylene glycol, 10% (w/v) PEG 8000. The successful crystallisation conditions, soaking conditions, and cryoprotectants used are detailed in Table S2.

### X-ray data collection and structure determination

Data were collected at Diamond Light Source (Didcot, UK) at 100 K using Pilatus 6M-F detectors and wavelengths of 0.92-0.98 Å. All data were processed using XDS (65). Further data processing and structural studies was carried out using CCP4 program package (66, 67). The apo structure of NagK was solved by the molecular replacement (MR) using the MR pipeline MORDA (68) with the best solution found for the model (PDB ID: 4DB3). The model was refined using REFMAC5 (69) and *PHENIX* (70) and rebuilt using COOT (71). The refined apo NagK model was used as a MR search model in MOLREP (72) for the NagK-GlcNAc-ADP data, which crystallised in a different space group. The MR solution was refined using *Buccaneer* (73), following which further refinement was performed as above. The crystals of NagK-GlcNAc-AMP, NagK-GlcNAc, NagK-GlcNAc-6’-phosphate and NagK-GlcNAc-AMP-PNP were in the same space group as the NagK-GlcNAc-ADP complex, however, phased MR (74) was used to reposition the small domain in the NagK-GlcNAc structure. All structures were subjected to phased refinement in REFMAC5 (75) with input DM phases (76) from NCS averaging. The models were validated using MOLPROBITY (77) implemented in the CCP4i2 interface (78).

### Molecular Dynamics

Molecular dynamics was performed in YASARA v.20.12.24 (79). The structure of NagK complexed with GlcNAc and AMP-PNP was cleaned to remove water and PEG molecules. Molecular dynamics was run using the md_runfast macro for 5 ns using the AMBER15FB force field (80). Simulations including divalent cations were performed by replacing water molecule 97 with the relevant cation.

## Acknowledgements

SR, MVV, and NJH were funded by BBSRC grant BB/N001591/1 and the UKRI CoA funds. JRA was funded by The National Science Foundation's Directorate of Human Resource Development’s Louis Stokes Alliance for Minority Participation Program Grant No. HRD-1408748, provided through the University of Oklahoma Office of Undergraduate Research.

## Author contributions

SR and NJH designed the study and NJH obtained funding. SR, MVV, JRA, NB, AK and KE collected data. SR, MNI and NJH analysed data. SR and NJH wrote the initial manuscript. All authors contributed to revision of the manuscript.

## Data availability

All data underpinning this work are publicly available. Structure coordinates and structure factor files are deposited with the Protein Data Bank (accession numbers: 7PA1, 7P7I, 7P7W, 7P9L, 7P9P and 7P9Y). Enzymatic and biophysical data are available as Supplementary Files or from Open Research Exeter (doi: to be confirmed on acceptance).

## References

1. Hascall V, Esko JD. Hyaluronan. In: rd, Varki A, Cummings RD, Esko JD, Stanley P, Hart GW, et al., editors. Essentials of Glycobiology. Cold Spring Harbor (NY)2015. p. 197–206.

2. Lindahl U, Couchman J, Kimata K, Esko JD. Proteoglycans and Sulfated Glycosaminoglycans. In: rd, Varki A, Cummings RD, Esko JD, Stanley P, Hart GW, et al., editors. Essentials of Glycobiology. Cold Spring Harbor (NY)2015. p. 207–21.

3. Tiemeyer M, Nakato H, Esko JD. Arthropoda. In: rd, Varki A, Cummings RD, Esko JD, Stanley P, Hart GW, et al., editors. Essentials of Glycobiology. Cold Spring Harbor (NY)2015. p. 335–49.

4. Zachara N, Akimoto Y, Hart GW. The O-GlcNAc Modification. In: rd, Varki A, Cummings RD, Esko JD, Stanley P, Hart GW, et al., editors. Essentials of Glycobiology. Cold Spring Harbor (NY)2015. p. 239–51.

5. Ogawa M, Okajima T. Structure and function of extracellular O-GlcNAc. Curr Opin Struct Biol. 2019;56:72–7.

6. Groves JA, Lee A, Yildirir G, Zachara NE. Dynamic O-GlcNAcylation and its roles in the cellular stress response and homeostasis. Cell Stress Chaperones. 2013;18(5):535–58.

7. Hardiville S, Hart GW. Nutrient regulation of signaling, transcription, and cell physiology by O-GlcNAcylation. Cell Metab. 2014;20(2):208–13.

8. Hart GW. Nutrient regulation of signaling and transcription. J Biol Chem. 2019;294(7):2211–31.

9. Seltmann G, Holst O. The bacterial cell wall. Berlin; New York: Springer; 2001. ix, 280 p. p.

10. Whitfield C, Trent MS. Biosynthesis and export of bacterial lipopolysaccharides. Annu Rev Biochem. 2014;83:99–128.

11. Reid AN, Szymanski CM. Biosynthesis and assembly of capsular polysaccharides. In: Holst O, Brennan PJ, von Itzstein M, editors. Microbial Glycobiology. London: Academic Press; 2010. p. 351–73.

12. Forsberg LS, Noel KD, Box J, Carlson RW. Genetic locus and structural characterization of the biochemical defect in the O-antigenic polysaccharide of the symbiotically deficient Rhizobium etli mutant, CE166. Replacement of N-acetylquinovosamine with its hexosyl-4-ulose precursor. J Biol Chem. 2003;278(51):51347–59.

13. Visansirikul S, Kolodziej SA, Demchenko AV. Staphylococcus aureus capsular polysaccharides: a structural and synthetic perspective. Org Biomol Chem. 2020;18(5):783–98.

14. Hong Y, Reeves PR. Diversity of o-antigen repeat unit structures can account for the substantial sequence variation of wzx translocases. J Bacteriol. 2014;196(9):1713–22.

15. Islam ST, Lam JS. Synthesis of bacterial polysaccharides via the Wzx/Wzy-dependent pathway. Can J Microbiol. 2014;60(11):697–716.

16. Hong Y, Morcilla VA, Liu MA, Russell EL, Reeves PR. Three Wzy polymerases are specific for particular forms of an internal linkage in otherwise identical O units. Microbiology. 2015;161(8):1639–47.

17. Lizak C, Gerber S, Numao S, Aebi M, Locher KP. X-ray structure of a bacterial oligosaccharyltransferase. Nature. 2011;474(7351):350–5.

18. Terra VS, Mills DC, Yates LE, Abouelhadid S, Cuccui J, Wren BW. Recent developments in bacterial protein glycan coupling technology and glycoconjugate vaccine design. J Med Microbiol. 2012;61(7):919–26.

19. Barreteau H, Kovac A, Boniface A, Sova M, Gobec S, Blanot D. Cytoplasmic steps of peptidoglycan biosynthesis. FEMS Microbiol Rev. 2008;32(2):168–207.

20. Milewski S, Gabriel I, Olchowy J. Enzymes of UDP-GlcNAc biosynthesis in yeast. Yeast. 2006;23(1):1–14.

21. Park JT. Identification of a dedicated recycling pathway for anhydro-N-acetylmuramic acid and N-acetylglucosamine derived from Escherichia coli cell wall murein. J Bacteriol. 2001;183(13):3842–7.

22. Plumbridge J. An alternative route for recycling of N-acetylglucosamine from peptidoglycan involves the N-acetylglucosamine phosphotransferase system in Escherichia coli. J Bacteriol. 2009;191(18):5641–7.

23. Popowska M, Osinska M, Rzeczkowska M. N-acetylglucosamine-6-phosphate deacetylase (NagA) of Listeria monocytogenes EGD, an essential enzyme for the metabolism and recycling of amino sugars. Arch Microbiol. 2012;194(4):255–68.

24. Yadav V, Panilaitis B, Shi H, Numuta K, Lee K, Kaplan DL. N-acetylglucosamine 6-phosphate deacetylase (nagA) is required for N-acetyl glucosamine assimilation in Gluconacetobacter xylinus. Plos One. 2011;6(6):e18099.

25. Fisher JF, Mobashery S. The sentinel role of peptidoglycan recycling in the beta-lactam resistance of the Gram-negative Enterobacteriaceae and Pseudomonas aeruginosa. Bioorg Chem. 2014;56:41–8.

26. Borisova M, Gaupp R, Duckworth A, Schneider A, Dalugge D, Muhleck M, et al. Peptidoglycan Recycling in Gram-Positive Bacteria Is Crucial for Survival in Stationary Phase. MBio. 2016;7(5).

27. Komatsuzawa H, Fujiwara T, Nishi H, Yamada S, Ohara M, McCallum N, et al. The gate controlling cell wall synthesis in Staphylococcus aureus. Mol Microbiol. 2004;53(4):1221–31.

28. Zhang W, Jones VC, Scherman MS, Mahapatra S, Crick D, Bhamidi S, et al. Expression, essentiality, and a microtiter plate assay for mycobacterial GlmU, the bifunctional glucosamine-1-phosphate acetyltransferase and N-acetylglucosamine-1-phosphate uridyltransferase. Int J Biochem Cell Biol. 2008;40(11):2560–71.

29. Oyeleye A, Normi Yahaya M. Chitinase: diversity, limitations, and trends in engineering for suitable applications. Bioscience Rep. 2018;38(4).

30. Rathore AS, Gupta RD. Chitinases from Bacteria to Human: Properties, Applications, and Future Perspectives. Enzyme Res. 2015;2015:791907.

31. Aggarwal C, Paul S, Tripathi V, Paul B, Khan MA. Chitinolytic activity in Serratia marcescens (strain SEN) and potency against different larval instars of Spodoptera litura with effect of sublethal doses on insect development. BioControl. 2015;60(5):631–40.

32. Meibom KL, Li XB, Nielsen AT, Wu CY, Roseman S, Schoolnik GK. The Vibrio cholerae chitin utilization program. Proc Natl Acad Sci U S A. 2004;101(8):2524–9.

33. Fang WX, Du T, Raimi OG, Hurtado-Guerrero R, Marino K, Ibrahim AFM, et al. Genetic and structural validation of Aspergillus fumigatus N-acetylphosphoglucosamine mutase as an antifungal target. Bioscience Rep. 2013;33:689–99.

34. Nishitani Y, Maruyama D, Nonaka T, Kita A, Fukami TA, Mio T, et al. Crystal structures of N-acetylglucosamine-phosphate mutase, a member of the alpha-D-phosphohexomutase superfamily, and its substrate and product complexes. Journal of Biological Chemistry. 2006;281(28):19740–7.

35. Dik DA, Fisher JF, Mobashery S. Cell-Wall Recycling of the Gram-Negative Bacteria and the Nexus to Antibiotic Resistance. Chem Rev. 2018;118(12):5952–84.

36. Weihofen WA, Berger M, Chen H, Saenger W, Hinderlich S. Structures of human N-acetylglucosamine kinase in two complexes with N-acetylglucosamine and with ADP/glucose: Insights into substrate specificity and regulation. J Mol Biol. 2006;364(3):388–99.

37. Roy S, Vega MV, Harmer NJ. Carbohydrate Kinases: A Conserved Mechanism Across Differing Folds. Catalysts. 2019;9(1).

38. Brigham CJ, Malamy MH. Characterization of the RokA and HexA broad-substrate-specificity hexokinases from *Bacteroides fragilis* and their role in hexose and *N*-acetylglucosamine utilization. J Bacteriol. 2005;187(3):890–901.

39. Conejo MS, Thompson SM, Miller BG. Evolutionary bases of carbohydrate recognition and substrate discrimination in the ROK protein family. J Mol Evol. 2010;70(6):545–56.

40. Hansen T, Schonheit P. ATP-dependent glucokinase from the hyperthermophilic bacterium *Thermotoga maritima* represents an extremely thermophilic ROK glucokinase with high substrate specificity. FEMS Microbiol Lett. 2003;226(2):405–11.

41. Coombes D, Davies JS, Newton-Vesty MC, Horne CR, Setty TG, Subramanian R, et al. The basis for non-canonical ROK family function in the N-acetylmannosamine kinase from the pathogen Staphylococcus aureus. J Biol Chem. 2020;295(10):3301–15.

42. Miyazono K, Tabei N, Morita S, Ohnishi Y, Horinouchi S, Tanokura M. Substrate recognition mechanism and substrate-dependent conformational changes of an ROK family glucokinase from *Streptomyces griseus*. J Bacteriol. 2012;194(3):607–16.

43. Nakamura T, Kashima Y, Mine S, Oku T, Uegaki K. Characterization and crystal structure of the thermophilic ROK hexokinase from *Thermus thermophilus*. J Biosci Bioeng. 2012;114(2):150–4.

44. Dorr C, Zaparty M, Tjaden B, Brinkmann H, Siebers B. The hexokinase of the hyperthermophile *Thermoproteus tenax*. ATP-dependent hexokinases and ADP-dependent glucokinases, two alternatives for glucose phosphorylation in Archaea. J Biol Chem. 2003;278(21):18744–53.

45. Hansen T, Reichstein B, Schmid R, Schonheit P. The first archaeal ATP-dependent glucokinase, from the hyperthermophilic crenarchaeon Aeropyrum pernix, represents a monomeric, extremely thermophilic ROK glucokinase with broad hexose specificity. J Bacteriol. 2002;184(21):5955–65.

46. Martinez J, Nguyen LD, Hinderlich S, Zimmer R, Tauberger E, Reutter W, et al. Crystal structures of N-acetylmannosamine kinase provide insights into enzyme activity and inhibition. J Biol Chem. 2012;287(17):13656–65.

47. Uehara T, Park JT. The N-acetyl-D-glucosamine kinase of Escherichia coli and its role in murein recycling. J Bacteriol. 2004;186(21):7273–9.

48. Brown JI, Page BDG, Frankel A. The application of differential scanning fluorimetry in exploring bisubstrate binding to protein arginine N-methyltransferase 1. Methods. 2019.

49. Llano-Sotelo B, Azucena EF, Jr., Kotra LP, Mobashery S, Chow CS. Aminoglycosides modified by resistance enzymes display diminished binding to the bacterial ribosomal aminoacyl-tRNA site. Chem Biol. 2002;9(4):455–63.

50. Bai N, Roder H, Dickson A, Karanicolas J. Isothermal Analysis of ThermoFluor Data can readily provide Quantitative Binding Affinities. Sci Rep. 2019;9(1):2650.

51. Minasov G, Wawrzak Z, Onopriyenko O, Skarina T, Papazisi L, Savchenko A, et al. 1.95 Angstrom Resolution Crystal Structure of N-acetyl-D-glucosamine kinase from Vibrio vulnificus. Protein Data Bank2012.

52. Pendleton JN, Gorman SP, Gilmore BF. Clinical relevance of the ESKAPE pathogens. Expert Rev Anti Infect Ther. 2013;11(3):297–308.

53. Shakir NA, Aslam M, Bibi T, Rashid N. ADP-dependent glucose/glucosamine kinase from Thermococcus kodakarensis: cloning and characterization. Int J Biol Macromol. 2021;173:168–79.

54. Szatmari D, Sarkany P, Kocsis B, Nagy T, Miseta A, Barko S, et al. Author Correction: Intracellular ion concentrations and cation-dependent remodelling of bacterial MreB assemblies. Sci Rep. 2020;10(1):18185.

55. Zeinert R, Martinez E, Schmitz J, Senn K, Usman B, Anantharaman V, et al. Structure-function analysis of manganese exporter proteins across bacteria. Journal of Biological Chemistry. 2018;293(15):5715–30.

56. Li J, Wang C, Wu Y, Wu M, Wang L, Wang Y, et al. Crystal structure of Sa239 reveals the structural basis for the activation of ribokinase by monovalent cations. J Struct Biol. 2012;177(2):578–82.

57. Nishimasu H, Fushinobu S, Shoun H, Wakagi T. Crystal structures of an ATP-dependent hexokinase with broad substrate specificity from the hyperthermophilic archaeon Sulfolobus tokodaii. J Biol Chem. 2007;282(13):9923–31.

58. Nocek B, Stein AJ, Jedrzejczak R, Cuff ME, Li H, Volkart L, et al. Structural studies of ROK fructokinase YdhR from Bacillus subtilis: insights into substrate binding and fructose specificity. J Mol Biol. 2011;406(2):325–42.

59. Bird LE. High throughput construction and small scale expression screening of multi-tag vectors in Escherichia coli. Methods. 2011;55(1):29–37.

60. Savitsky P, Bray J, Cooper CD, Marsden BD, Mahajan P, Burgess-Brown NA, et al. High-throughput production of human proteins for crystallization: the SGC experience. J Struct Biol. 2010;172(1):3–13.

61. Peranen J, Rikkonen M, Hyvonen M, Kaariainen L. T7 vectors with modified T7lac promoter for expression of proteins in Escherichia coli. Anal Biochem. 1996;236(2):371–3.

62. Cook PF, Cleland WW. Enzyme kinetics and mechanism. London; New York: Garland Science; 2007. xxii, 404 p. p.

63. Cornish-Bowden A. Fundamentals of enzyme kinetics. 4th, completely revised and greatly enlarged edition. ed. Weinheim, Germany: Wiley-Blackwell; 2012. xviii, 498 pages p.

64. Harmer NJ, Vivoli Vega M. Reaction Chemical Kinetics in Biology. Biomolecular and Bioanalytical Techniques2019. p. 179–217.

65. Kabsch W. Xds. Acta Crystallogr D. 2010;66:125–32.

66. Potterton E, Briggs P, Turkenburg M, Dodson E. A graphical user interface to the CCP4 program suite. Acta Crystallogr D Biol Crystallogr. 2003;59(Pt 7):1131–7.

67. Winn MD, Ballard CC, Cowtan KD, Dodson EJ, Emsley P, Evans PR, et al. Overview of the CCP4 suite and current developments. Acta Crystallogr D. 2011;67:235–42.

68. Vagin A, Lebedev A. *MoRDa*, an automatic molecular replacement pipeline. Acta Crystallographica Section A - Foundations and Devices. 2015;A71:s19.

69. Murshudov GN, Skubak P, Lebedev AA, Pannu NS, Steiner RA, Nicholls RA, et al. REFMAC5 for the refinement of macromolecular crystal structures. Acta Crystallogr D. 2011;67:355–67.

70. Afonine PV, Grosse-Kunstleve RW, Echols N, Headd JJ, Moriarty NW, Mustyakimov M, et al. Towards automated crystallographic structure refinement with phenix.refine. Acta Crystallographica Section D-Structural Biology. 2012;68:352–67.

71. Emsley P, Lohkamp B, Scott WG, Cowtan K. Features and development of Coot. Acta Crystallogr D. 2010;66:486–501.

72. Vagin A, Teplyakov A. Molecular replacement with MOLREP. Acta Crystallogr D. 2010;66:22–5.

73. Cowtan K. Completion of autobuilt protein models using a database of protein fragments. Acta Crystallogr D Biol Crystallogr. 2012;68(Pt 4):328–35.

74. Vagin AA, Isupov MN. Spherically averaged phased translation function and its application to the search for molecules and fragments in electron-density maps. Acta Crystallographica Section D-Structural Biology. 2001;57:1451–6.

75. Pannu NS, Murshudov GN, Dodson EJ, Read RJ. Incorporation of prior phase information strengthens maximum-likelihood structure refinement. Acta Crystallogr D. 1998;54:1285–94.

76. Cowtan K. Recent developments in classical density modification. Acta Crystallogr D. 2010;66:470–8.

77. Williams CJ, Headd JJ, Moriarty NW, Prisant MG, Videau LL, Deis LN, et al. MolProbity: More and better reference data for improved all-atom structure validation. Protein Sci. 2018;27(1):293–315.

78. Potterton L, Agirre J, Ballard C, Cowtan K, Dodson E, Evans PR, et al. CCP4i2: the new graphical user interface to the CCP4 program suite. Acta Crystallographica Section D-Structural Biology. 2018;74:68–84.

79. Krieger E, Vriend G. New ways to boost molecular dynamics simulations. J Comput Chem. 2015;36(13):996–1007.

80. Wang LP, Martinez TJ, Pande VS. Building Force Fields: An Automatic, Systematic, and Reproducible Approach. J Phys Chem Lett. 2014;5(11):1885–91.

81. Laskowski RA, Swindells MB. LigPlot+: multiple ligand-protein interaction diagrams for drug discovery. J Chem Inf Model. 2011;51(10):2778–86.

82. Wallace AC, Laskowski RA, Thornton JM. LIGPLOT: a program to generate schematic diagrams of protein-ligand interactions. Protein Eng. 1995;8(2):127–34.

