## Supplementary information for "Structure and function of *N*-acetylglucosamine kinase illuminates the catalytic mechanism of ROK kinases"

Roy et al.

Supplementary information

**Figure S1: Purification of NagK from diverse organisms**. Proteins were expressed in *E. coli* as 6-His-SUMO (*V. vulnificus* and *Plesiomonas shigelloides*) and 6-His-GST fusion proteins. Proteins were purified using immobilised metal affinity chromatography and size exclusion chromatography (SEC) using an ÄKTAxpress system. Samples from size exclusion were analysed by SDS-PAGE to verify the identity and purity of the proteins. Samples were mixed 1:1 with loading buffer and run on ExpressPlus 4-20% gels (Genscript # M42015) using the manufacturer's supplied MOPS buffer at 150 V for 50 minutes and stained with InstantBlue (Abcam #ab119211). Samples: M: 5 μL Spectra Multicolor Broad Range Protein Ladder (Thermo Scientific # 26623); C: 2 μL resuspended cells; P; 2 μL post-lysis pellet; S: 2 μL post-lysis supernatant; FT: 3 μL IMAC flow-through; SEC samples: 10 μL from samples at the SEC peak. **A**: 6-His-SUMO-NagK from *P. shigelloides*. **B**: 6_His-GST-NagK from *Photobacterium damselae*. **C**: 6-His-GST-NagK from *Pseudoalteromonas* sp. P1-8. **D**: 6-His-SUMO-NagK from *Vibrio vulnificus*.

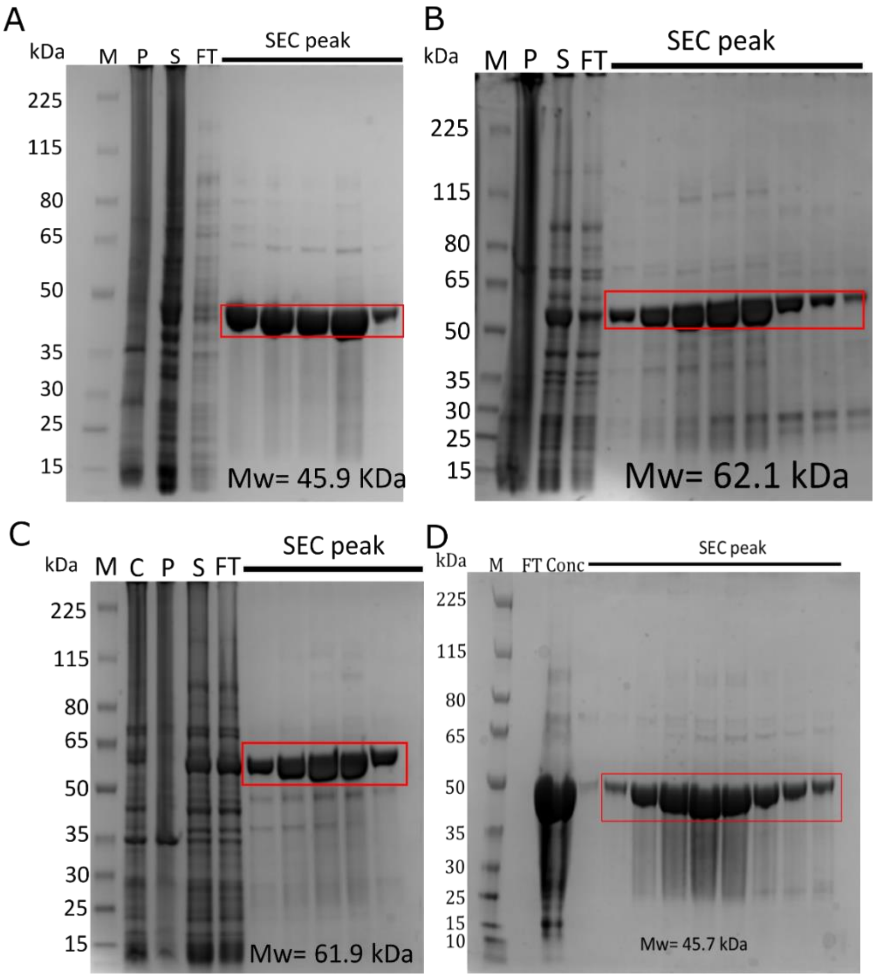

**Figure S2: Verification of NagK protein identity**. Purified NagK from the experiments above was separated by SDS-PAGE and western blot. The blot was probed using an iBind (Thermo Scientific) following the manufacturer's instructions with mouse anti-penta-His (Qiagen # 34650) primary and IRDye® 680RD Goat anti-Mouse (LI-COR # 926-68070) secondary antibodies used at 1:2000 and 1:5000 dilutions respectively.

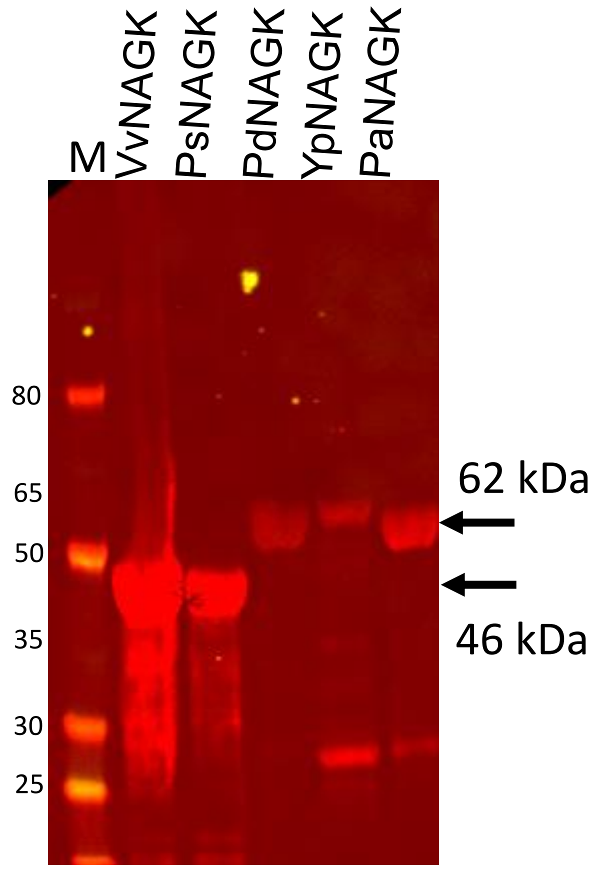

**Figure S3: NagK products are weak inhibitors.** Product inhibition of NagK by GlcNAc-6-phosphate and ADP were assayed. Inhibition by GlcNAc-6-phosphate was tested using the same assay as used for GlcNAc activity, using NagK at 180 ng/mL, GlcNAc at 100 μM and ATP at 300 μM; ADP inhibition was measured using a coupled assay with the enzymes *N*-acetylglucosamine-phosphate mutase (NagP; from *Candida albicans*; (1)), *N*-acetylglucosamine-1-phosphate uridyltransferase (GlmU; from *Escherichia coli*; (2)), and UDP-N-acetylgalactosamine dehydrogenase (WbpO; also has activity against UDP-GlcNAc; from *Pseudomonas aeruginosa* serotype O6; (3)). Each enzyme was purified in the same manner as described for NagK. The assay mixture contained 40 mM Hepes pH 7.5, 100 mM KCl, 8 mM $MgCl_2$, 50 mM $NH_4(SO_4)_2$, 5 mM DTT, 100 μg/mL BSA, 300 μM ATP, 100 μM GlcNAc, 200 μM UTP, 5 mM $NAD^+$, 1 μg/mL NagK, 20 μg/mL NagP, 5 μg/mL GlmU, and 20 μg/mL WbpO. The reaction was followed at 340 nm for 2000 s and the rate determined following the lag phase. Data were fitted to the revised Morrison Ki equation (4). The Morrison Ki for GlcNAc-6-phosphate was determined as 40 ± 4 mM, whilst the determined Ki for ADP was 4.7 ± 0.4 mM.

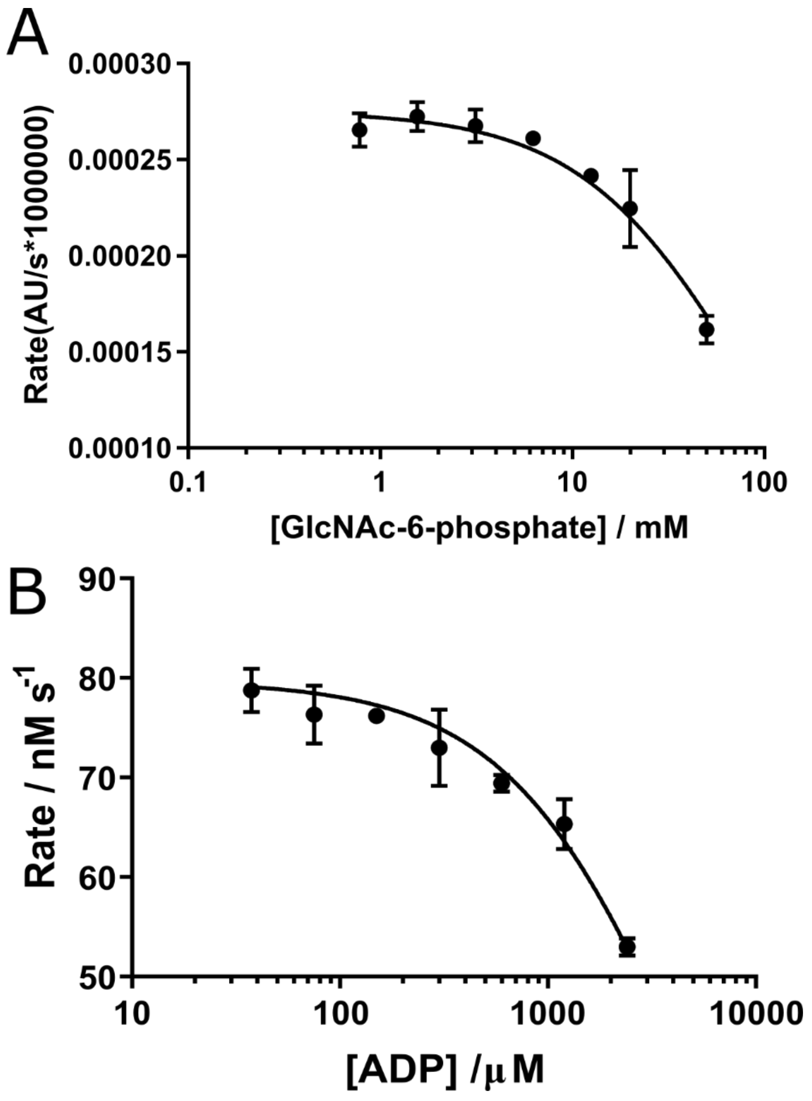

**Figure S4: Binding of ATP to generate a ternary complex requires a further rotation of the small domain.** Upon binding of AMP-PNP (spheres, carbon atoms coloured white), the small domain (left) rotates relative to the large domain. There is a 16° rotation from the position of the GlcNAc complex (magenta, GlcNAc carbon atoms coloured yellow; PDB ID: 7P9Y) to the GlcNAc-AMP-PNP complex (orange; PDB ID: 7P9P). Structures were superimposed over the large domain. Atom colours: nitrogen, blue; oxygen, yellow; chloride, green, potassium, purple. Figure generated using PyMOL v. 2.4.1 (5).

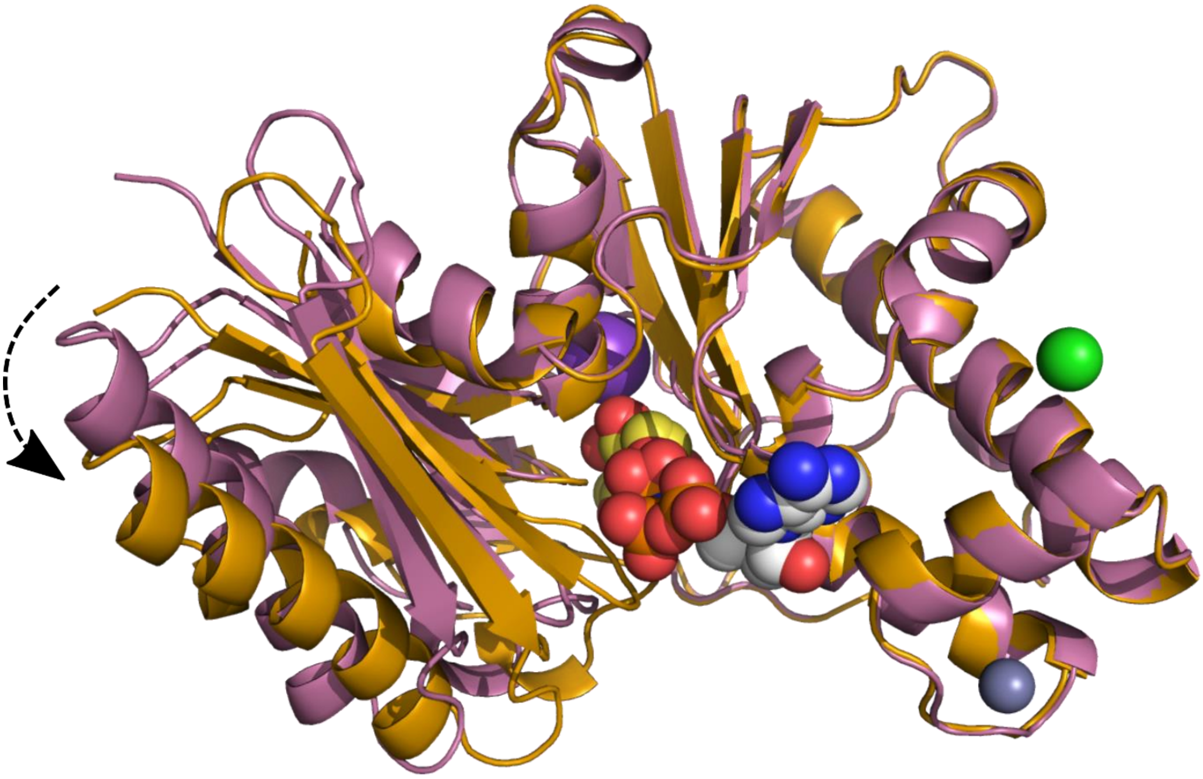

**Figure S5: Sequence alignment of NagK orthologues.** The sequences chosen share 39-78% sequence identity. Possible active site residues are indicated: dark blue arrow, proposed metal binding site (D6); orange box, ATP phosphate binding region (G9-K11); orange arrows, side chain ATP binding (T132, E196); sky blue arrow, main chain ATP binding (G255); green arrows, GlcNAc binding (S78, N104, E154, H157, D187); black arrow, proposed catalytic base (D105). Figure produced using ESpript v.3.0 (6).

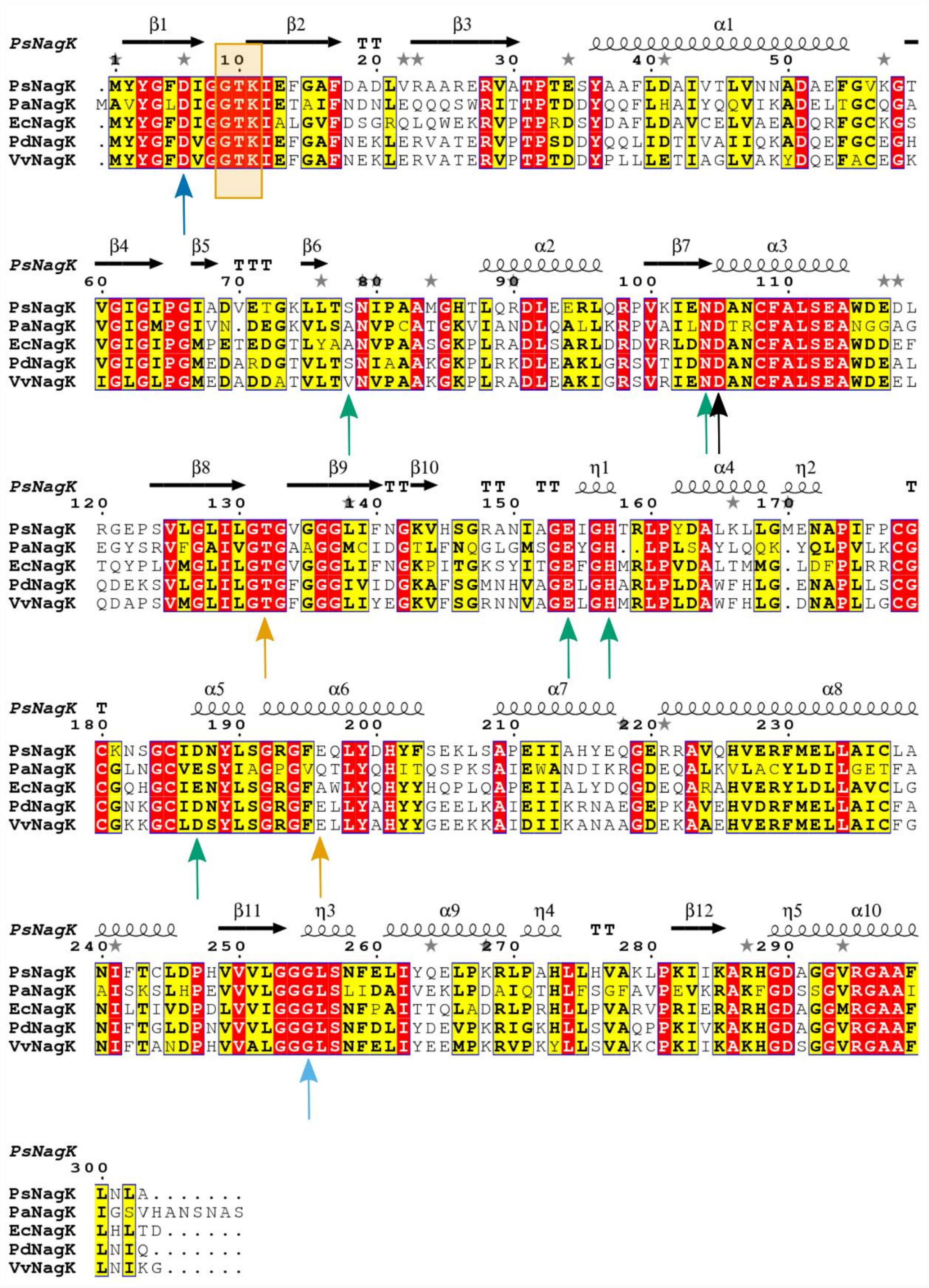

**Figure S6: The NagK ternary complex with GlcNAc and AMP-PNP shows the same conformation as a previous ROK kinase ternary complex including the catalytic metal.** The structure of NagK in complex with AMP-PNP (black dashed arrow) and GlcNAc (black solid arrow) (**A**; PDB ID: 7P9P) was superimposed with the structure of human NanK in complex with ADP (grey dashed arrow), ManNAc (grey solid arrow) and magnesium (black arrowhead) (**C**; PDB ID: 2YHY) using Pymol v2.5, focusing on the larger domain (right in this image). The superposition (**B**) shows that the conformation of the small domain relative to the large domain is well conserved between the two structures. The magnesium position is consistent with this being the catalytic cation position in the NagK structure. Colours: *Ps*NagK: green; *Hs*NanK: sky blue; nitrogen atoms: blue; oxygen atoms: red; phosphorus atoms: orange spheres; zinc ions: grey spheres; magnesium/calcium ions: light green spheres; potassium ions: purple spheres. Images generated using Pymol v 2.5 (5).

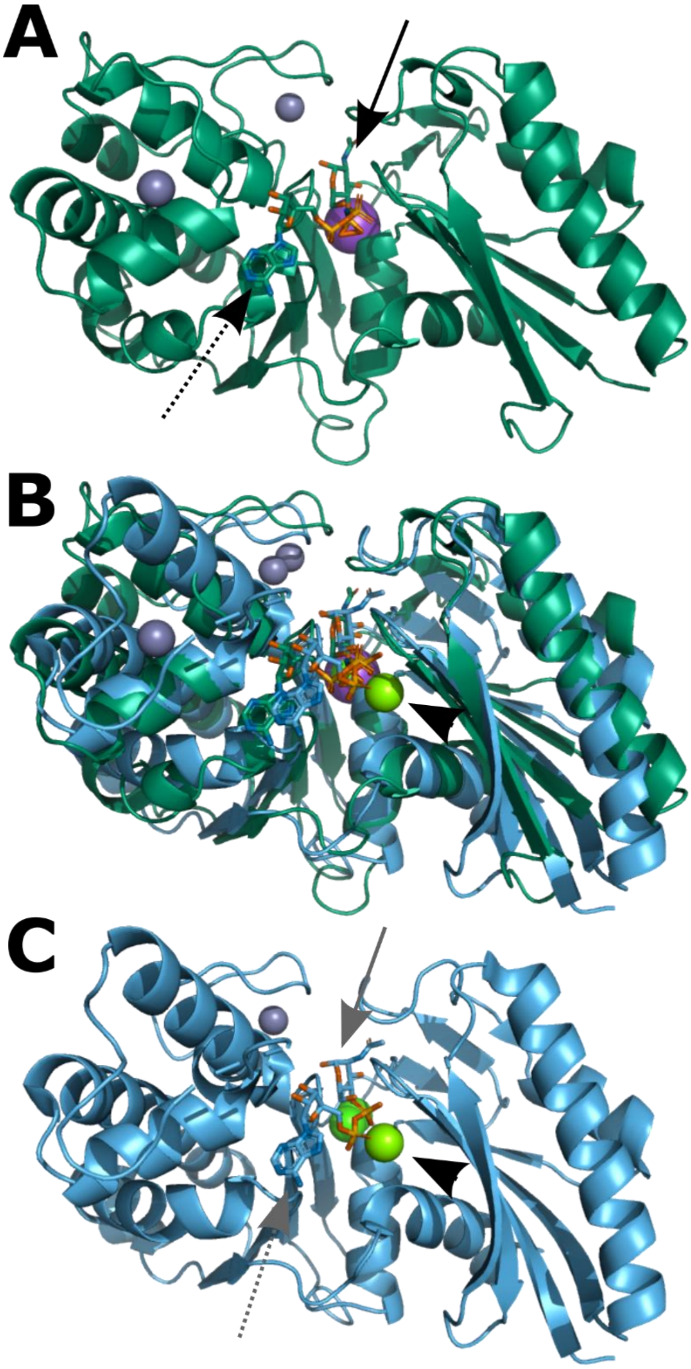

**Figure S7: Molecular dynamics confirms the likely metal binding site.** The NagK-GlcNAc-AMP-PNP ternary complex (PDB ID: 7P9P) was altered to replace the nitrogen in AMP-PNP with oxygen (A). In separate structures, a bound water between D6 and AMP-PNP was replaced with magnesium (B), calcium (C) or manganese (D). Molecular dynamics was run for 5 ns using YASARA v.20.12.24. Images show the proposed metal binding site, with amino acids within 4.5 Å of the metal shown as sticks and the remainder of NagK as cartoon. Figure generated using PyMOL v. 2.4.1 (5).

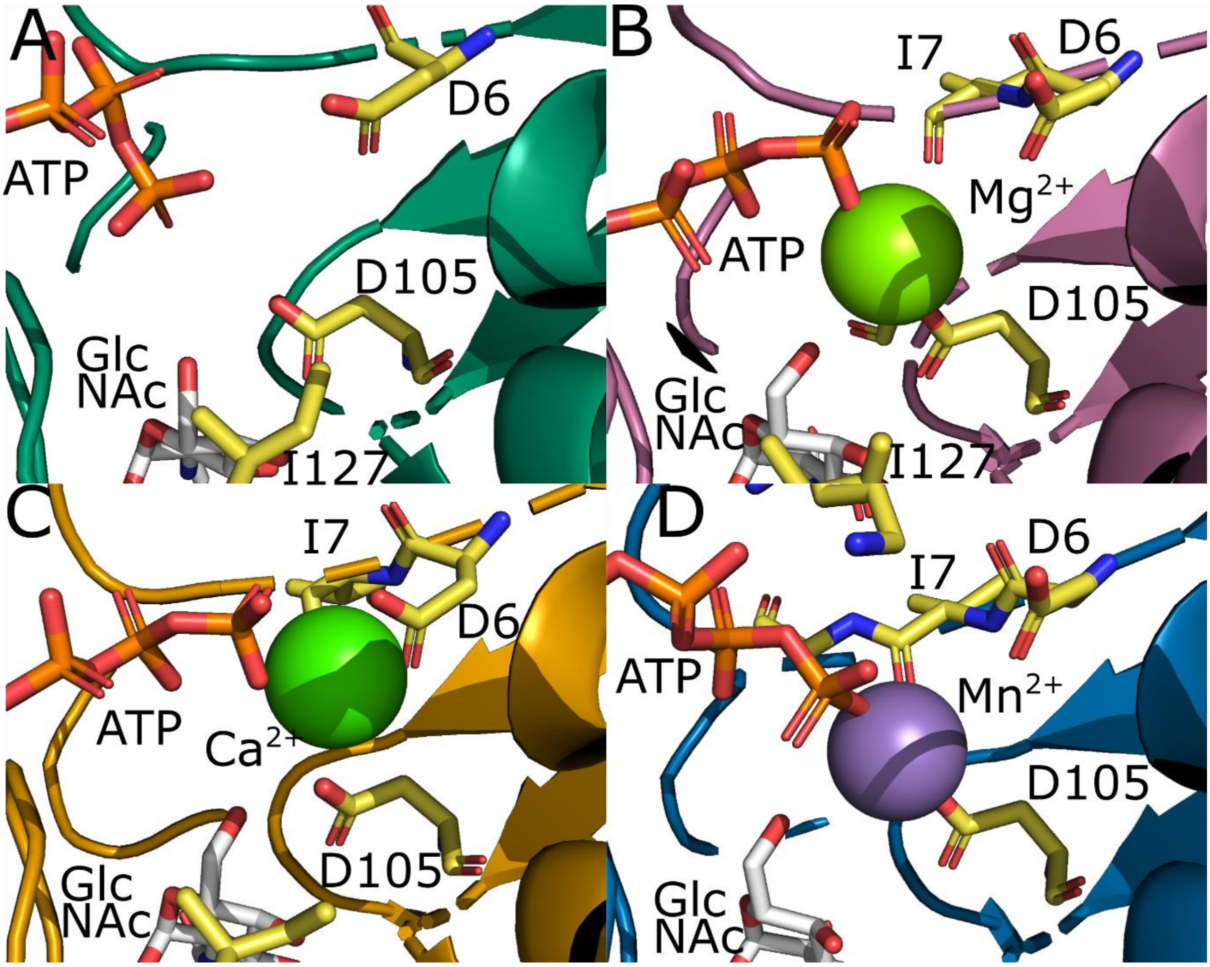

**Figure S8: Sequence alignment of ROK kinases.** Sequences of six ROK kinases representing the major known activities were aligned, using available crystal structures. 3OHR: fructokinase from *Bacillus subtilis* (7); 2AA4: *N*-acetylmannosamine kinase from *E. coli*; 3HTV: D-allose kinase from *E. coli*. 3VGL: glucokinase from *Streptomyces griseus* (8); 1WOQ: polyphosphate glucokinase from *Arthrobacter* sp. KM (9). NagK proposed active site residues are indicated: dark blue arrow, proposed metal binding site (D6); orange box, ATP phosphate binding region (G9-K11); orange arrows, side chain ATP binding (T132, E196); sky blue arrow, main chain ATP binding (G255); green arrows, GlcNAc binding (S78, N104, E154, H157, D187); black arrow, proposed catalytic base (D105). Figure produced using ESpript v.3.0 (6).

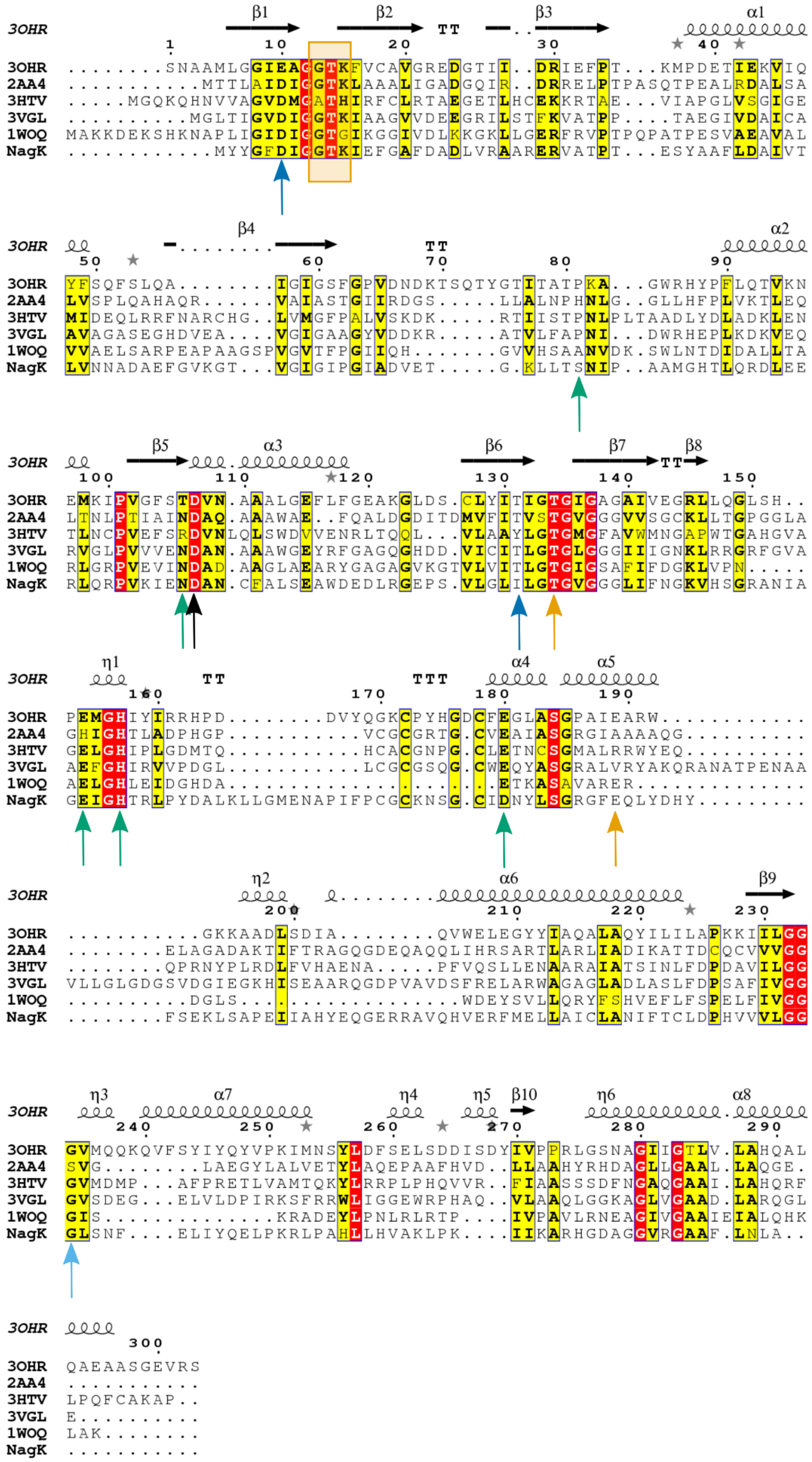
3OHR β1 β2 TT β3 α1
1 10 20 30 40
3OHR ........SNAAMLGGIEAGGTKFVCAVGREDGTII..DRIEFPT...KMPDETIEKVIQ
2AA4 ..........MTTLAIDIGGTKLAAALIGADGQIR..DRRELPTPASQTPEALRDALSA
3HTV ....MGQKQHNVVAGVDMGATHIRFCLRTAEGETLHCEKKRTAE...VIAPGLVSGIGE
3VGL .........MGLTIGVDIGGTKIAAGVVDEEGRILSTFKVATPP....TAEGIVDAICA
1WOQ MAKKDEKSHKNAPLIGIDIGGTGIKGGIVDLKKGKLLGERFRVPTPQPATPESVAEAVAL
NagK ...........MYYGFDIGGTKIEFGAFDADLVRAARERVATPT...ESYAAFLDAIVT
3OHR β4 TT α2
50 60 70 80 90
3OHR YFSQFSLQA........IGIGSFGPVDNDKTSQTYGTITATPKA...GWRHYPFLQTVKN
2AA4 LVSPLQAHAQR......VAIASTGIIRDGS.....LLALNPHNLG..GLLHFPLVKTLEQ
3HTV MIDEQLRRFNARCHG..LVMGFPALVSKDK.....RTIISTPNLPLTAADLYDLADKLEN
3VGL AVAGASEGHDVEA....VGIGAAGYVDDKR.....ATVLFAPNI...DWRHEPLKDKVEQ
1WOQ VVAELSARPEAPAAGSPVGVTFPGIIQH.......GVVHSAANVDK.SWLNTDIDALLTA
NagK LVNNADAEFGVKGT...VGIGIPGIADVET.....G.KLLTSNIP..AAMGHTLQRDLEE
3OHR β5 α3 β6 β7 TT β8
100 110 120 130 140 150
3OHR EMKIPVGFSTDVN.AALGEFLFGEAKGLDS.CLYITIGTGIGAGAIVEGRLLQGLSH..
2AA4 LTNLPTIAINDAQ.AAAWAE..FQALDGDITDMVFITVSTGVGGGVVSGCKLLTGPGGLA
3HTV TLNCPVEFSRDVNLQLSWDVVENRLTQQL...VLAAYLGTGMGFAVWMNGAPWTGAHGVA
3VGL RVGLPVVVENDAN.AAAWGEYRFGAGQGHDD.VICITLGTGLGGGIIIGNKLRRGRFGVA
1WOQ RLGRPVEVINDAD.AAGLAEARYGAGAGVKGTVLVITLGTGIGSAFIFDGKLVPN.....
NagK RLQRPVKIENDAN.CFALSEAWDEDLRGEPS.VLGLILGTGVGGGLIFNGKVHSGRANIA
3OHR η1 TT TTT α4 α5
160 170 180 190
3OHR PEMGHIYIRRHPD........DVYQGKCPYHGDCFEGLASGPAIEARW............
2AA4 GHIGHTLADPHGP...........VCGCGRTG.CVEAIASGRGIAAAAQG..........
3HTV GELGHIPLGDMTQ...........HCACGNPG.CLETNCSGMALRRWYEQ..........
3VGL AEFGHIRVVPDGL...........LCGCGSQG.CWEQYASGRALVRYAKQRANATPENAA
1WOQ AELGHLEIDGHDA......................ETKASAVARER..............
NagK GEIGHTRLPYDALKLLGMENAPIFPCGCKNSG.CIDNYLSGRGFEQLYDHY.........
3OHR η2 α6 β9
200 210 220 230
3OHR ..........GKKAADLSDIA........QVWELEGYYIAQALAQYILILAPKKIILGG
2AA4 ........ELAGADAKTIFTRAGQGDEQAQQLIHRSARTLARLIADIKATTDCQCVVVGG
3HTV ........QPRNYPLRDLFVHAENA....PFVQSLLENAARAIATSINLFDPDAVILGG
3VGL VLLGLGDGSVDGIEGKHISEAARQGDPVAVDSFRELARWAGAGLADLASLFDPSAFIVGG
1WOQ .........DGLS...................WDEYSVLLQRYFSHVEFLFSPELFIVGG
NagK ........FSEKLSAPEIIAHYEQGERRAVQHVERFMELLAICLANIFTCLDPHVVLGG
3OHR η3 α7 η4 η5 β10 η6 α8
240 250 260 270 280 290
3OHR GVMQQKQVFSYIYQYVPKIMNSYLDFSELSDDISDYIVPPRLGSNAGIIGTLV.LAHQAL
2AA4 SVG........LAEGYLALVETYLAQEPAAFHVD..LLAAHYRHDAGLLGAAL.LAQGE.
3HTV GVMDMP...AFPRETLVAMTQKYLRRPLPHQVVR..FIAASSSDFNGAQGAAI.LAHQRF
3VGL GVSDEG...ELVLDPIRKSFRRWLIGGEWRPHAQ..VLAAQLGGKAGLVGAAD.LARQGL
1WOQ GIS.............KRADEYLPNLRLRTP....IVPAVLRNEAGIVGAAIEIALQHK
NagK GLSNF....ELIYQELPKRLPAHLLHVAKLPK....IIKARHGDAGGVRGAAF.LNLA..
3OHR
300
3OHR QAEAASGEVRS
2AA4 ...........
3HTV LPQFCAKAP..
3VGL E..........
1WOQ LAK........
NagK ...........

**Table S1: Crystal information, date collection and refinement:**

| Project | NagK | NagK - GlcNAc | NagK- GlcNAc -ADP | NagK – GlcNAc AMP-PNP | NagK- GlcNAc - P | NagK – AMP-PNP |
|---|---|---|---|---|---|---|
| **Data collection statistics** | | | | | | |
| **Beamline** | I04 Diamond | I03 Diamond | I04-1 Diamond | I04 Diamond | I03 Diamond | I03 Diamond |
| **Wavelength (Å)** | 0.9795 | 0.9763 | 0.9159 | 0.9795 | 0.9763 | 0.9763 |
| **Space group** | $P3_121$ | $P3_221$ | $P3_221$ | $P3_221$ | $P3_221$ | $P6_5$ |
| **Unit Cell Parameters a, b, c (Å)** | 95.2, 95.2, 180.6 | 115.5, 115.5, 119.7.3 | 115.1, 115.1, 120.3 | 115.2, 115.2, 120.4 | 114.5, 114.5, 119.3 | 121.3, 121.3, 91.7 |
| **α, β, γ (°)** | 90.0, 90.0, 120.0 | 90.0, 90.0 120.0 | 90.0, 90.0, 120.0 | 90.0, 90.0, 120.0 | 90.0, 90.0, 120.0 | 90.0, 90.0, 120.0 |
| **Resolution range (Å)[a]** | 75.01 – 1.70 (1.73-1.70) | 57.74 – 1.94 (1.99 – 1.94) | 99.68 – 1.57 (1.60 - 1.57) | 57.60 – 2.11 (2.17 – 2.11) | 99.17-1.75 (1.78 - 1.75) | 60.63 – 2.20 (2.27 – 2.20) |
| **Total reflections [a]** | 521,556 (25,890) | 457,524 (29,499) | 1,273,708 (54,820) | 535,990 (43,641) | 346,458 (17,705) | 127,050 (10,922) |
| **Unique reflections [a]** | 104,319 (5,073) | 68,568 (4,548) | 128,365 (6,335) | 53,533 (4,354) | 90,389 (4,505) | 38,637 (3,361) |
| **Completeness (%)[a]** | 99.6 (99.7) | 100.0 (100.0) | 100.0 (100.0) | 100.0 (100.0) | 99.1, (100.0) | 99.2 (99.8) |
| **Multiplicity [a]** | 5.0 (5.1) | 6.7 (6.5) | 9.9 (8.7) | 10.0 (10.0) | 3.8 (3.9) | 3.3 (3.2) |
| **$R_{merge}$ (%) [a,b]** | 4.6 (262.5) | 8.6 (231.3) | 10.1 (321.7) | 17.3 (336.0) | 9.1 (169.1) | 17.8 (272.5) |
| **$\langle I\rangle/\langle\sigma(I)\rangle$ [a]** | 13.8 (0.4) | 11.0 (0.8) | 12.7 (0.6) | 9.5 (0.7) | 8.4 (0.8) | 5.2 (0.4) |
| **$CC_{1/2}$ [a,c]** | 0.999 (0.270) | 0.999 (0.282) | 0.999 (0.291) | 0.998 (0.336) | 0.998 (0.194) | 0.964 (0.396) |
| **Wilson B-factor [d] (Å$^2$)** | 41.3 | 48.5 | 33.4 | 49.9 | 35.1 | 48.9 |
| **Refinement statistics** | | | | | | |
| **$R_{work}$** | 0.183 | 0.208 | 0.185 | 0.191 | 0.194 | 20.9 |
| **$R_{free}$** | 0.210 | 0.250 | 0.212 | 0.233 | 0.223 | 25.2 |
| **No. of protein monomers** | 2 | 2 | 2 | 2 | 2 | 2 |

| in a.u. | | | | | | |
|---|---|---|---|---|---|---|
| **Number of atoms** | | | | | | |
| **Macromolecules** | 4,891 | 4,679 | 4,976 | 4,741 | 4,899 | 4,666 |
| **Ligands and metal ions** | 2 | 34 | 86 | 96 | 42 | 64 |
| **Solvent** | 597 | 382 | 749 | 360 | 511 | 242 |
| **Number of protein residues** | 609 | 608 | 610 | 610 | 609 | 608 |
| **RMS bond lengths (Å)** | 0.009 | 0.009 | 0.007 | 0.007 | 0.012 | 0.006 |
| **RMS bond angles (°)** | 1.62 | 1.67 | 1.46 | 1.57 | 1.77 | 1.42 |
| **Ramachandran favored (%)[e]** | 97.9 | 94.4 | 98.0 | 96.2 | 97.9 | 96.2 |
| **Ramachandran outliers (%)[e]** | 0.0 | 0.2 | 0.0 | 0.0 | 0.0 | 0.5 |
| **Clashscore [e]** | 4.05 | 12.6 | 8.8 | 6.9 | 7.0 | 6.3 |
| **Average B-factor protein (Å$^2$)** | 42.2 | 55.3 | 27.8 | 49.3 | 35.1 | 46.5 |
| **Average B-factor ligands (Å$^2$)** | 39.2 | 69.1 | 30.3 | 56.8 | 40.3 | 72.7 |
| **Average B-factor solvent (Å$^2$)** | 55.4 | 56.9 | 43.5 | 57.3 | 49.6 | 50.7 |
| **RCBS PDB code** | 7P7I | 7P9Y | 7P7W | 7P9P | 7P9L | 7PA1 |

[a]Values for the highest resolution shell are given in parentheses

[b] $R_{merge} = \Sigma_h \Sigma_i |I_{h,i} - \langle I_h \rangle| / \Sigma_h \Sigma_i I_{h,I}$

[c] $CC_{1/2}$ is defined in (10)

[d] Wilson B-factor was estimated by SFCHECK (11)

[e] The Ramachandran statistics and clashscore were calculated using MOLPROBITY (12).

**Table S2: Crystallisation conditions**

| NagK/NagK-substrate complex | Morpheus screen position/conditions | Co-crystallised | Soaking substrate | Soaking/cryoprotection solution | Soaking time (sec) |
|---|---|---|---|---|---|
| NagK (Native) | A12 (60 mM divalent cations; 0.1 M Tris/bicine pH 8.5; 12.5% each MPD, PEG 1K, PEG 3350 | N/A | N/A | 25% (v/v) MPD, 20% (w/w) PEG 1K, 20% (w/w) PEG 3350, 50 mM Tris pH 8.5 | N/A |
| NagK with GlcNAc | B8 (90 mM halogens; 0.1 M Na-HEPES/MOPS pH 7.5; 12.5% each MPD, PEG 1K, PEG 3350) | 500 µM GlcNAc | N/A | 25% (v/v) MPD, 20% (w/w) PEG 1K, 20% (w/w) PEG 3350, 50 mM MOPS pH 7.5 | N/A |
| NagK with GlcNAc and AMP | D7 (120 mM alcohols; 0.1 M Na-HEPES/MOPS pH 7.5; 30% each glycerol and PEG 4K) | 500 µM GlcNAc | 10 mM AMP | 30% (v/v) glycerol, 15% (w/w) PEG 4K, 100 mM MOPS pH 7.5 | 30 |
| NagK with GlcNAc and AMP-PNP | A3 (60 mM divalent cations; 0.1 M imidazole/MES pH 6.5; 30% each glycerol and PEG 4K) | 500 µM GlcNAc | 10 mM AMP-PNP | 30% (v/v) glycerol, 15% (w/w) PEG 4K, 100 mM imidazole pH 6.5 | 90 |
| NagK with GlcNAc and ADP | D3 (120 mM alcohols; 0.1 M imidazole/ MES pH 6.5; 30% each glycerol and PEG 4K) | 500 µM GlcNAc | 10 mM ADP | 30% (v/v) glycerol, 15% (w/w) PEG 4K, 100 mM imidazole pH 6.5 | 60 |
| NagK with GlcNAc-6-phosphate | B9 (90 mM halogens; 0.1 M Tris/bicine pH 8.5; 30% each PEG 550 MME and PEG 20K) | 10 mM GlcNAc-6-phosphate and 500 µM ATP | N/A | 35% PEG 500-MME, 15% PEG 8000, 100 mM Tris pH 8.5, 10 mM ATP | 60 |
| NagK with AMP-PNP | C9 (0.09M NPS; 0.1 M Tris/bicine pH 8.5; 30% each PEG 550 MME and PEG 20K) | N/A | 10 mM AMP-PNP | 30% (v/v) PEG400, 15% (w/w) PEG 4K, 100 mM Tris pH 8.5 | 90 |

**References:**


1. Nishitani, Y., Maruyama, D., Nonaka, T., Kita, A., Fukami, T. A., Mio, T., Yamada-Okabe, H., Yamada-Okabe, T., and Miki, K. (2006) Crystal structures of N-acetylglucosamine-phosphate mutase, a member of the alpha-D-phosphohexomutase superfamily, and its substrate and product complexes. *J Biol Chem* **281**, 19740-19747
2. Olsen, L. R., Vetting, M. W., and Roderick, S. L. (2007) Structure of the E. coli bifunctional GlmU acetyltransferase active site with substrates and products. *Protein Sci* **16**, 1230-1235
3. Zhao, X., Creuzenet, C., Belanger, M., Egbosimba, E., Li, J., and Lam, J. S. (2000) WbpO, a UDP-N-acetyl-D-galactosamine dehydrogenase from Pseudomonas aeruginosa serotype O6. *J Biol Chem* **275**, 33252-33259
4. Cook, P. F., and Cleland, W. W. (2007) *Enzyme kinetics and mechanism*, Garland Science, London ; New York
5. The PyMOL Molecular Graphics System. 2.4.1 Ed., Schrödinger, LLC.
6. Robert, X., and Gouet, P. (2014) Deciphering key features in protein structures with the new ENDscript server. *Nucleic Acids Res* **42**, W320-324
7. Nocek, B., Stein, A. J., Jedrzejczak, R., Cuff, M. E., Li, H., Volkart, L., and Joachimiak, A. (2011) Structural studies of ROK fructokinase YdhR from Bacillus subtilis: insights into substrate binding and fructose specificity. *J Mol Biol* **406**, 325-342
8. Miyazono, K., Tabei, N., Morita, S., Ohnishi, Y., Horinouchi, S., and Tanokura, M. (2012) Substrate recognition mechanism and substrate-dependent conformational changes of an ROK family glucokinase from *Streptomyces griseus*. *J Bacteriol* **194**, 607-616
9. Mukai, T., Kawai, S., Mori, S., Mikami, B., and Murata, K. (2004) Crystal structure of bacterial inorganic polyphosphate/ATP-glucomannokinase. Insights into kinase evolution. *J Biol Chem* **279**, 50591-50600
10. Karplus, P. A., and Diederichs, K. (2012) Linking crystallographic model and data quality. *Science* **336**, 1030-1033
11. Vaguine, A. A., Richelle, J., and Wodak, S. J. (1999) SFCHECK: a unified set of procedures for evaluating the quality of macromolecular structure-factor data and their agreement with the atomic model. *Acta Crystallogr D Biol Crystallogr* **55**, 191-205
12. Williams, C. J., Headd, J. J., Moriarty, N. W., Prisant, M. G., Videau, L. L., Deis, L. N., Verma, V., Keedy, D. A., Hintze, B. J., Chen, V. B., Jain, S., Lewis, S. M., Arendall, W. B., 3rd, Snoeyink, J., Adams, P. D., Lovell, S. C., Richardson, J. S., and Richardson, D. C. (2018) MolProbity: More and better reference data for improved all-atom structure validation. *Protein Sci* **27**, 293-315
